# Dysregulated Airway Host Defense in Hyper IgE Syndrome due to STAT3 Mutations

**DOI:** 10.1101/2024.08.14.607930

**Authors:** Ling Sun, Samantha A. Walls, Hong Dang, Nancy L. Quinney, Patrick R. Sears, Taraneh Sadritabrizi, Koichi Hasegawa, Kenichi Okuda, Takanori Asakura, Xiuya Chang, Meiqi Zheng, Yu Mikami, Felicia U. Dizmond, Daniela Danilova, Lynn Zhou, Anshulika Deshmukh, Deborah M. Cholon, Giorgia Radicioni, Troy D. Rogers, William J. Kissner, Matthew R. Markovetz, Tara N. Guhr Lee, Mark I. Gutay, Charles R. Esther, Michael Chua, Barbara R. Grubb, Camille Ehre, Mehmet Kesimer, David B. Hill, Lawrence E. Ostrowski, Brian Button, Martina Gentzsch, Chevalia Robinson, Kenneth N. Olivier, Alexandra F. Freeman, Scott H. Randell, Wanda K. O’Neal, Richard C. Boucher, Gang Chen

## Abstract

**Rationale:** Hyper IgE syndrome (STAT3-HIES), also known as Job’s syndrome, is a rare immunodeficiency disease typically caused by dominant-negative STAT3 mutations. STAT3-HIES syndrome is characterized by chronic pulmonary infection and inflammation, suggesting impairment of pulmonary innate host defense.

**Objectives:** To identify airway epithelial host defense defects consequent to STAT3 mutations that, in addition to reported mutant STAT3 immunologic abnormalities, produce pulmonary infection.

**Methods:** STAT3-HIES sputum was evaluated for biochemical/biophysical properties. STAT3-HIES excised lungs were harvested for histology; bronchial brush samples were collected for RNA sequencing and in vitro culture. A STAT3-HIES-specific mutation (R382W), expressed by lentiviruses, and a STAT3 knockout, generated by CRISPR/Cas9, were maintained in normal human bronchial epithelia under basal or inflammatory (IL1β) conditions. Effects of STAT3 deficiency on transcriptomics, and epithelial ion channel, secretory, antimicrobial, and ciliary functions were assessed.

**Measurements and Main Results:** Mucus concentrations and viscoelasticity were increased in STAT3-HIES sputum. STAT3-HIES excised lungs exhibited mucus obstruction and elevated IL1β expression. STAT3 deficiency impaired CFTR-dependent fluid and mucin secretion, inhibited expression of antimicrobial peptides, cytokines, and chemokines, and acidified airway surface liquid at baseline and post-IL1β exposure in vitro. Notably, mutant STAT3 suppressed IL1R1 expression. STAT3 mutations also inhibited ciliogenesis in vivo and impaired mucociliary transport in vitro, a process mediated via HES6 suppression. Administration of a γ-secretase inhibitor increased HES6 expression and improved ciliogenesis in STAT3 R382W mutant cells.

**Conclusions:** STAT3 dysfunction leads to multi-component defects in airway epithelial innate defense, which, in conjunction with STAT3-HIES immune deficiency, contributes to chronic pulmonary infection.

## Introduction

STAT3-HIES is a rare immunodeficiency disease usually caused by dominant-negative STAT3 mutations (1, 2) that produce deficiency of STAT3 functions (3). Impaired STAT3 activity has been reported to produce immune cell dysfunctions that impede bacterial and fungal host defense (4). While STAT3-HIES is largely considered a primary immunodeficiency, evidence suggests that compromise of innate defense in non-hematopoietic cells also contributes to STAT3-HIES pulmonary (5) and intestinal pathologies (6). For example, systemic immune cell functions were largely restored by hematopoietic stem cell transplantation in STAT3-HIES patients (7) and Stat3-mutant mice (8). However, resistance to infection, epithelial antimicrobial peptide secretion, and tissue remodeling activities were not completely restored in the lung or gut (7, 8).

Mucociliary clearance (MCC) is a major innate airway epithelial host defense activity that mediates clearance of infectious agents in lung (9). MCC requires the coordinated function of multiple epithelial cell types, including secretory and ciliated cells. The cystic fibrosis transmembrane conductance regulator (CFTR) and mucin 5B (MUC5B) are predominantly expressed by secretory cells (10), but CFTR is also expressed in rare ionocytes (10–13). In cystic fibrosis (CF), defective CFTR limits fluid, but not mucin, secretion, producing mucus hyperconcentration, impaired MCC, and infection (14). The airway mucus of STAT3-HIES has been described as thick and tenacious (15, 16), suggesting that STAT3 mutations may also disrupt the balance of fluid vs mucin secretion (17) in STAT3-HIES airways.

Defective STAT3 has also been reported to impair ciliogenesis and ciliated cell repair (5, 18, 19). Cilia not only provide the motive force to propel pathogens and inhaled particles trapped in the mucus layer from the lung but also mechanically sense and auto-regulate mucus concentration via an extracellular ATP-mediated pathway (20). Thus, dysregulated ciliary function may also contribute to tenacious STAT3-HIES mucus, similar to other genetic lung diseases, e.g., primary ciliary dyskinesia (PCD) (17).

In this study, we tested the hypothesis that STAT3 regulates multiple components of innate airway epithelial defense. These studies employed sputum and bronchial epithelial samples from people with STAT3-HIES and primary human bronchial epithelial (HBE) cells modified to express a patient-specific STAT3 mutation R382W (2) or loss of STAT3 by CRISPR/Cas9. The roles of STAT3 in the coordinated regulation of airway epithelial MCC elements (mucus concentrations, cilia numbers/function) and bactericidal activities, including regulation of antimicrobial peptide (AMP) secretion and airway surface liquid (ASL) pH, were investigated.

## Methods

### Patient information

Healthy controls and people with STAT3-HIES (Supplementary table 1) were recruited to the NIH Clinical Center, and specimens were collected following IRB-approved protocols. Detailed methods are available in the Online Supplement. All bioinformatic data generated from this study were deposited in GEO.

## Results

### STAT3-HIES lungs exhibit mucus hyperconcentration and IL1β-mediated inflammation

STAT3-HIES sputum exhibited higher mucus concentrations, as indexed by %solids (21), than controls, comparable to CF levels (14) (Figure 1A). STAT3-HIES sputum also exhibited elevated biomarkers of mucin concentration, including sialic acid mass and sialic acid-to-urea ratios (Figure 1B), and a trend for total mucin proteins (Figure 1C) (22). STAT3-HIES sputum contained both MUC5AC and MUC5B mucins (Figures 1D), and a disproportionate increase in MUC5AC (23), resulting in an increased MUC5AC:MUC5B ratio (Figure 1E). As predicted by concentration (24), STAT3-HIES sputum mucus complex viscosity was raised (Figure 1F).

**Figure 1:**
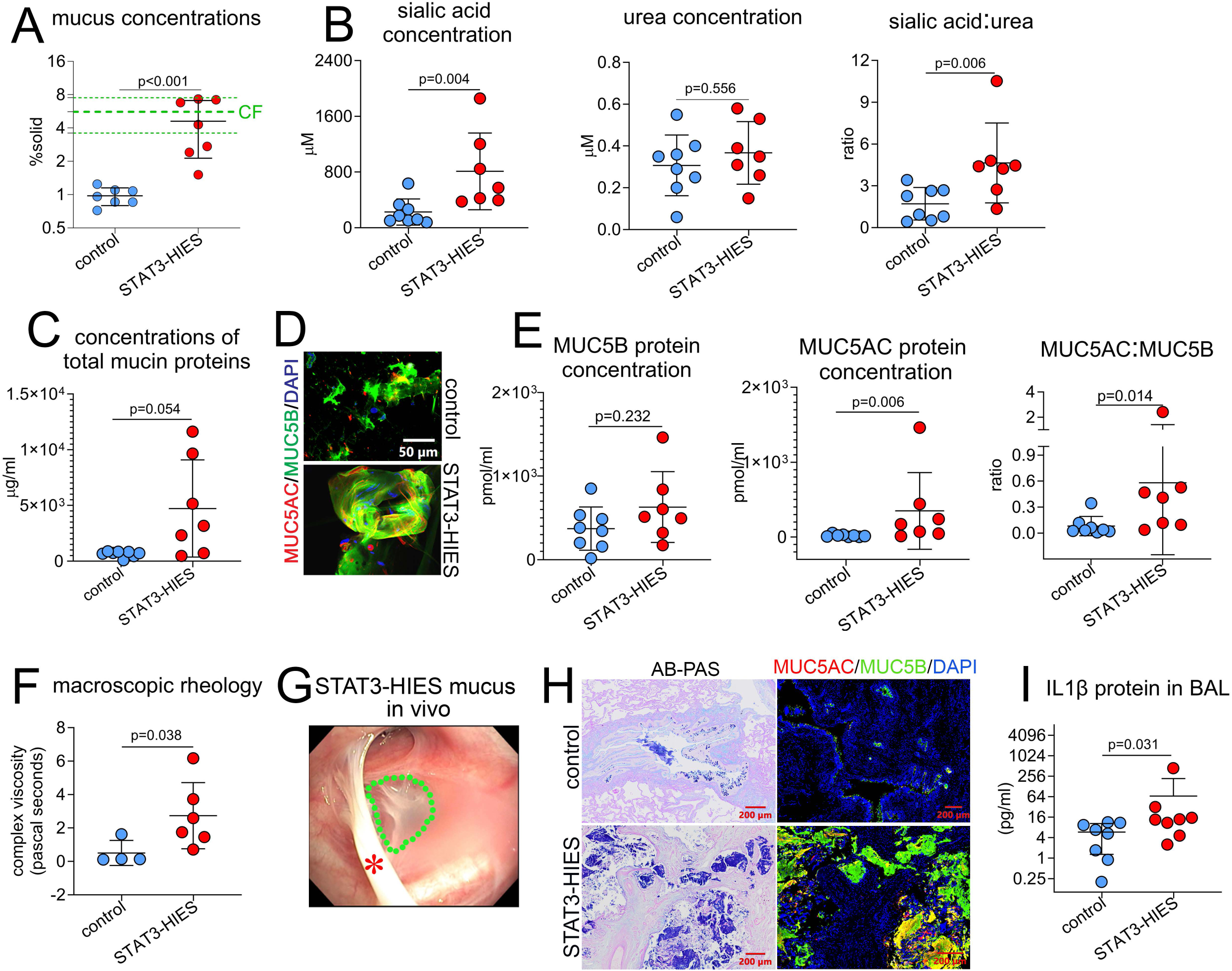
STAT3-HIES lung exhibits muco-obstructive inflammation. **(A)** Sputum specimens were collected from control and STAT3-HIES donors (n=7/group). Mucus concentrations were calculated by measuring the dry to wet ratio of sputum. The range of CF mucus % solids from a previous study was presented as mean ± SD (thick and thin green dash lines, respectively). **(B)** Sialic acid and urea concentrations, and their ratios in sputum were measured by mass spectrometry and analyzed. **(C)** Total mucin concentrations in sputum were measured with the use of size-exclusion chromatography and differential refractometry. **(D)** Expression of MUC5B and MUC5AC proteins in sputum was detected by IF staining. **(E)** MUC5B and MUC5AC concentrations, and their ratios in sputum were measured by internal standard labeled mass spectrometry and analyzed. **(F)** Complex viscosity of sputum was measured by shear dependence of the viscosity by rheometer. **(G)** Viscous mucus overlaying bronchus of right upper lobe of a STAT3-HIES individual was imaged during bronchoscopy. Red asterisk indicates mucus strand, green dash line circles mucus plaque presented in the bronchus. **(H)** AB-PAS and MUC5AC/MUC5B IF staining were performed in the distal airways in control and STAT3-HIES excised lung tissues. **(I)** IL1β concentration in BAL of control and STAT3-HIES lungs (n=8/group) was determined by mesoscale U-Plex platform. IF: immunofluorescence, BAL: bronchoalveolar lavage. AB-PAS: Alcian blue-periodic acid Schiff. Histology images in (D, H) are representatives of n=3/group.

Bronchoscopy and autopsy lungs were utilized to study STAT3-HIES mucus in vivo. Bronchoscopy visualized mucus plaques and strands on bronchial surfaces of people with STAT3-HIES (Figure 1G). Mucus plugging was visualized in distal airways of autopsy lungs, as shown by AB-PAS and MUC5B/5AC immunofluorescence (IF) staining (Figure 1H, Supplementary Figure 1A). IL1β promotes mucus hyperconcentration in CF and non-CF bronchiectasis (25, 26), and IL1β expression was increased in STAT3-HIES lung tissues compared to controls as well as in STAT3-HIES bronchoalveolar lavage (BAL) samples (Supplementary Figure 1B, Figure 1I).

### STAT3 mutations inhibit CFTR expression and function

To identify mechanisms relating STAT3 mutations to hyperconcentrated mucus, bulk RNA-seq analyses were performed on bronchial brushing tissue from STAT3-HIES and control subjects. The gene expression profiles formed genotype-distinctive clusters by principal component analysis (Suppl Fig.2A). Since STAT3-HIES shared hyperconcentrated mucus phenotype with CF, we first examined CFTR expression. CFTR mRNA was decreased in STAT3-HIES, whereas expression of markers of major CFTR expressing cell types [SCGB1A1, SCGB3A1 and KRT7 for secretory cells (10, 27); FOXI1 for ionocytes] were not (Figure 2A, Suppl Fig.2B,C). Expression of other ion channel genes regulating ASL hydration, e.g., via Cl^-^ channel (ANO1, SLC26A9) (28, 29) or the subunits of epithelial Na^+^ channel (ENaC, including SCNN1A, B) (30), were similar, and SCNN1G was decreased, in STAT3-HIES compared to controls. (Suppl Fig.2D).

**Figure 2:**
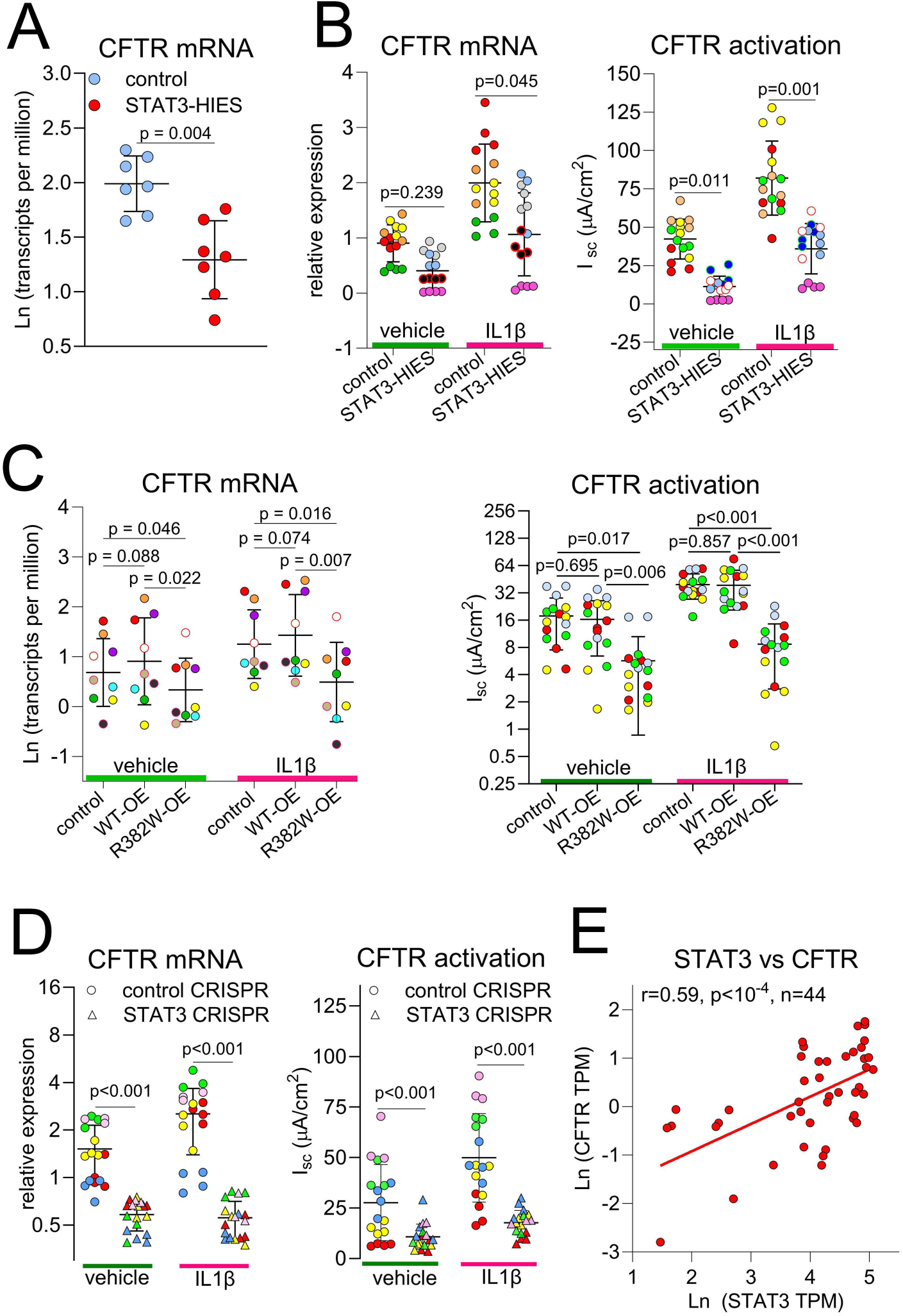
STAT3 deficiency leads to reduced CFTR expression and activity. **(A)** Total RNA was freshly isolated from bronchial brushing tissues of healthy control and STAT3-HIES donors (7 donors/group), and CFTR mRNA was measured by bulk RNA-seq. **(B)** Primary HBE cells isolated from control and STAT3-HIES bronchial brushing tissues (4 donors/group) were expanded and differentiated under ALI conditions prior to IL1β exposure. CFTR mRNA and activity were measured by TaqMan assays and Ussing chamber assay, respectively. **(C)** Normal HBE cells (n=9 donors) were transduced with lentiviruses expressing control vector, wild type STAT3 (WT-OE) and mutant STAT3 (R382W-OE) and cultured under ALI prior to IL1β exposure followed by bulk RNA-seq. CFTR activity in response to forskolin were measured and quantified. **(D)** Normal HBE cells (n=5 donors) were electroporated with control or STAT3 CRISPR/Cas9, followed by ALI culture prior to IL1β exposure. CFTR mRNA and activity of each culture were quantified. **(E)** Linear regression test was performed to STAT3 and CFTR mRNAs of well-differentiated normal HBE cells from 44 donors (measured by RNA-seq), the Ln transformed Pearson correlation coefficient (r) was reported in the graph. Symbols with same colors in the same panel indicate multiple cultures originated from the same donor except in (**A**, **E**). ALI: air-liquid interface; HBE: human bronchial epithelial; OE: overexpression; Ln: natural logarithm; TMP: transcripts per million.

To test whether decreased CFTR expression in bronchial brushing specimens produced functional defects in Cl^-^/fluid transport, primary HBE cells from STAT3-HIES and control bronchial brushing samples were cultured under air-liquid interface (ALI) conditions. To mimic the regulation of CFTR and mucin associated with chronic inflammation in STAT3-HIES lung in vivo, cultures were studied under basal and IL1β-treated conditions. In both vehicle and IL1β treated cultures, CFTR mRNA and protein expression trended lower in STAT3-HIES HBE cells compared to controls (Figure 2B, Suppl. Fig. 2E). Importantly, the acute activation of CFTR function (forskolin response) was reduced under both conditions in STAT3-HIES HBE cells (Figure 2B). Consistent with bronchial brush data, expression of secretory cell and ionocyte markers, frequency of secretory cells, and mRNA for ion channels, e.g., TMEM16A (ANO1) and the ENaC subunits, were comparable between STAT3-HIES and control HBE cultures, regardless of IL1β treatment (Suppl Fig.2F-H). Collectively, these data suggest STAT3 deficiency leads to mucus hyperconcentration at least partially via reduction of CFTR function and not via altered differentiation of secretory cells, ionocytes, or expression of other non-CFTR channels known for ion/fluid secretion/absorption.

### STAT3 R382W mutation produces dysfunctional phospho-STAT3 signaling

Lentiviruses overexpressing wild-type STAT3 (WT-OE), the STAT3 R382W mutant (R382W-OE), a hotspot mutation in STAT3-HIES individuals (2), and empty lentiviral vector (control), were generated (Suppl Fig.3A). Both WT-OE and R382W-OE transduced HBE cells exhibited elevated STAT3 protein levels compared to controls after 1 and 4 weeks at ALI conditions (Suppl Fig.3B). However, phospho-STAT3 (p-STAT3) nuclear localization was reduced in R382W-OE transduced HBE cells at baseline and after IL1β exposure (Suppl Fig.3C) consistent with the previous report (31).

To quantify R382W-OE dysregulation of p-STAT3 induction and nuclear translocation, p-STAT3 responses to IL6 (a classic STAT3 activation cytokine) were tested using the human bronchial epithelial cell line H441. R382W-OE did not prevent STAT3 from phosphorylation. However, it decreased the fold induction of p-STAT3 and its nuclear translocation in response to IL-6 compared to WT-OE (Suppl Fig. 3D-F), providing a candidate mechanism linking dominant-negative mutations to impaired transcriptional activity due to defective nuclear localization of STAT3 R382W in HBE cells. *<u>STAT3 mutation or ablation inhibits CFTR expression and function.</u>* R382W-OE HBE cells exhibited a reduction in CFTR mRNA, and a trend of decreased CFTR protein compared to WT-OE or control at baseline and after IL1β exposure. (Figure 2C, Suppl Fig.3G). Similar to primary STAT3-HIES cells, CFTR mediated Cl^-^ transport was reduced in R382W-OE cells under both conditions (Figure 2C). Secretory cell markers/frequencies were moderately increased and FOXI1 mRNA levels were not changed, in both conditions in R382W-OE HBE cells (Suppl Fig.3H). These data suggest that R382W-OE failed to activate CFTR expression/function likely due to STAT3-mediated transcriptional dysregulation on a per-cell basis.

To confirm the role of STAT3 to regulate CFTR, a STAT3 knock-out (KO) model was generated in normal HBE cells by CRISPR/Cas9 (Suppl Fig.4A,B). STAT3-KO reduced CFTR mRNA, protein, and activation at baseline and/or in responses to IL1β (Figure 2D, Suppl Fig.4C). Further, compared to control lentivirus, expression of STAT3 WT-OE lentivirus in primary STAT3-HIES HBE cells increased CFTR expression (Suppl Fig.4D). A moderate positive correlation (32) between STAT3 and CFTR mRNAs (r=0.59, p<10^-4^) was detected in normal well-differentiated HBE cultures by bulk RNA-seq (Figure 2E), supporting a transcriptional activator role for STAT3 to regulate CFTR expression in HBE cells.

### STAT3 mediates an IL1β-induced GP130/JAK1 pathway to regulate CFTR expression

The IL6/ STAT3 axis regulates airway epithelial differentiation/regeneration (18, 19). We speculated that IL1β induced IL6 secretion from HBE (33) cells to activate STAT3 and CFTR. Indeed, IL1β exposure increased IL6 mRNA and protein within 24 hours (Suppl Fig.5A, Figure 3A). Induction of p-STAT3 by IL1β was evident after 8hrs, preceding the peak of CFTR mRNA induction that occurred at 24hrs post-IL1β exposure in normal HBE cells (Figure 3B). To block IL6 signaling, the IL6R inhibitor Tocilizumab (TCZ) was administered to normal HBE cells. TCZ strongly inhibited IL6-induced p-STAT3, but it only marginally reduced IL1β-induced p-STAT3 and did not decrease IL1β-induced CFTR mRNA (Figure 3C, Suppl Fig.5B). Further, IL6 itself did not increase CFTR mRNA (Suppl Fig.5C). We conclude that IL1β-induced STAT3 activation and CFTR expression are independent of the IL6/STAT3 cascade.

**Figure 3:**
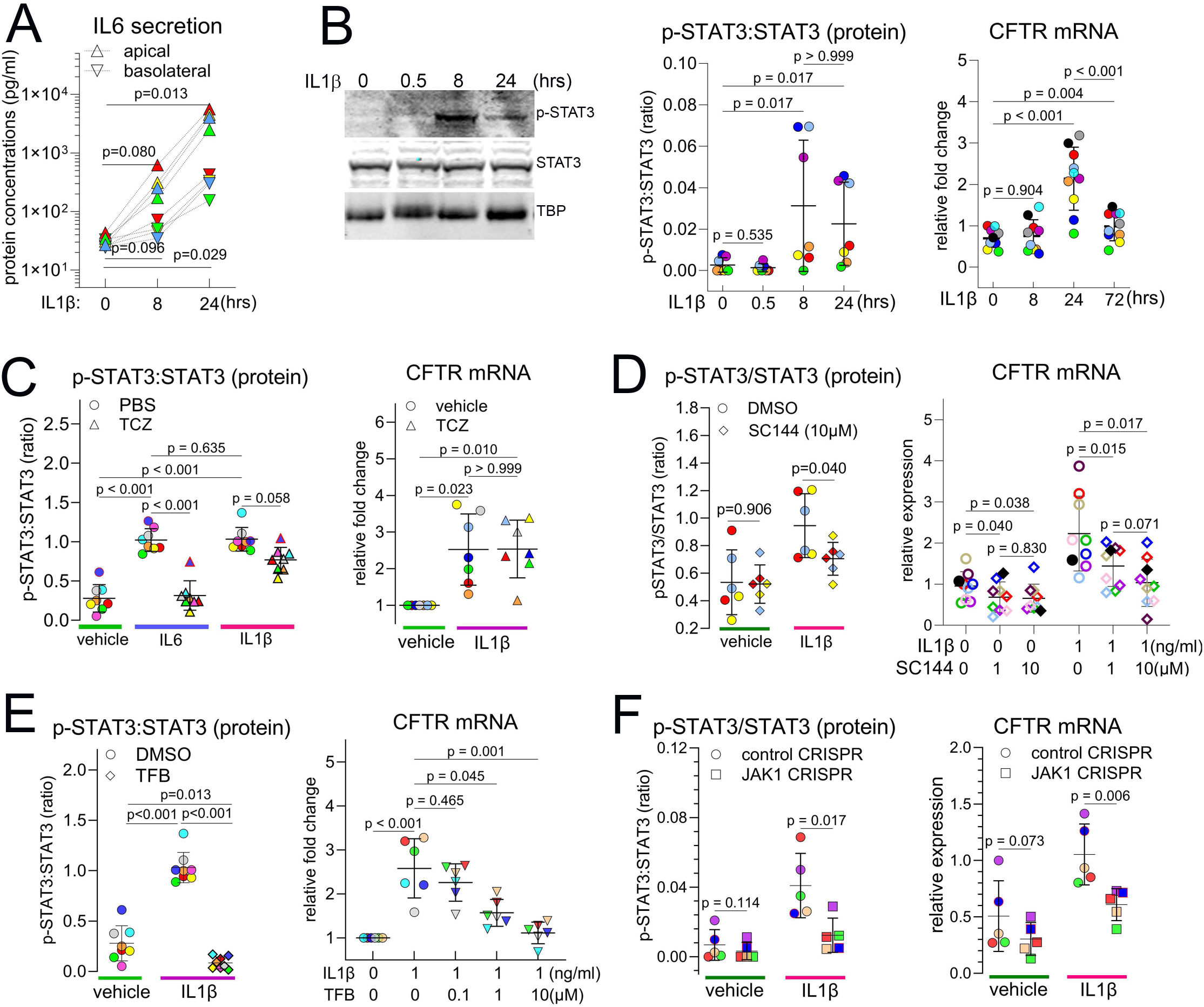
GP130-JAK1-STAT3 axis mediates IL1β-induced STAT3-phosphorylation and CFTR expression in normal HBE cells. **(A)** Normal HBE cells (n=4 donors) were exposed to IL1β, and apical and basolateral secretion of IL6 was measured by ELISA at 0, 8 and 24 hours after treatment. **(B)** Expression of p-STAT3 and STAT3 proteins were detected by western blot and their ratios were calculated in the normal HBE cells (n=7 donors) exposed with IL1β for 0, 8 and 24 hours, and CFTR mRNA (n=10 donors) was measured up to 72 hours. **(C)** To block IL6 signaling, Tocilizumab (TCZ) was administered to normal HBE cells (n=8 donors) prior to exposure to IL6 or IL1β. p-STAT3 and STAT3 proteins were detected, and their ratios were calculated. The CFTR mRNA fold changes in response to IL1β with or without TCZ was detected by TaqMan assays (n=7 donors). **(D)** To inhibit GP130 functions, SC144 was administered to the normal HBE cells (n=3 donors) prior to exposure to IL1β; p-STAT3 and STAT3 proteins were detected and their ratios were calculated. The CFTR mRNA change in response to IL1β was measure in normal HBE cells (n=9 donors) treated with SC144 at 0 (DMSO), 1 and 10 µM. **(E)** To inhibit JAK activity, Tofacitinib (TFB, 1µM) was administered to normal HBE cells (n=8 donors) prior to exposure to IL1β. The ratio of p-STAT3 to STAT3 proteins and CFTR mRNA (n=6 donors) was measured. **(F)** Normal HBE cells (n=5 donors) were electroporated with control or JAK1 CRISPR/Cas9, followed by ALI culture prior to exposure to IL1β. The ratio of p-STAT3 to STAT3 was calculated after western blot and densitometry quantification and CFTR mRNA was measured. Symbols with the same colors in the same panel indicate multiple cultures originated from the same donor, regardless of shapes. ALI: air-liquid interface; HBE: human bronchial epithelial.

STAT proteins can alternately be activated via the GP130/JAKs axis (34). To test whether GP130 and JAK mediate IL1β-induced p-STAT3 and CFTR expression, specific GP130 (SC144) and JAK (Tofacitinib, TFB) inhibitors were tested (35, 36). Administration of SC144, TFB or CRISPR/Cas9 targeting JAK1 (the dominant JAK isoform in HBE) inhibited IL1β-induced p-STAT3 and CFTR expression (Figure 3D-F, Suppl Fig.5D-H). We conclude that the GP130/JAK1 pathway mediates IL1β-induced p-STAT3 and CFTR in airway epithelia.

### STAT3 is required for coordinated mucin and fluid secretion

Compared to CF, STAT3-HIES exhibits reduced, but not complete loss of, CFTR function. However, STAT3-HIES subjects exhibit CF-like mucus hyperconcentration/plugging (Figure 1A) and recurrent infection, suggesting mutant STAT3 impairs MCC components beyond CFTR. The effects of STAT3 mutations on MUC5B and MUC5AC transcription and secretion were accordingly investigated. Despite higher mucin concentrations in sputum (Figure 1C), MUC5B and MUC5AC mRNAs were not increased in STAT3-HIES bronchial brushing specimens (Figure 4A) or cultured STAT3-HIES HBE cells (Suppl Fig.6A). Moreover, STAT3 R382W-OE and KO was associated with decreased MUC5B/MUC5AC mRNA levels and protein secretion in HBE cells under basal and/or IL1β-stimulated conditions, consistent with previous reports on mucin gene regulation by STAT3 (37, 38) (Figure 4B,C and Suppl Fig.6B,C). Expression of the MUC5B-interacting protein BPIFB1 that regulates MCC and mucus viscosity (39) was also decreased in STAT3-deficient HBE cells (Suppl Fig.6D), as were mucin-associated transcription factors, XBP1S (Figure 4D), SPDEF and FOXA3 (40) (Suppl Fig.6E). As predicted by the requirement for coordination of epithelial mucin and fluid secretion, MUC5B and MUC5AC mRNAs were tightly correlated with CFTR mRNA in response to IL1β in normal HBE cells (Suppl Fig.6F). These correlations were enhanced in STAT3 WT-OE, but lost in R382W-OE HBE cells, in response to IL1β (Figure 4E,F).

**Figure 4:**
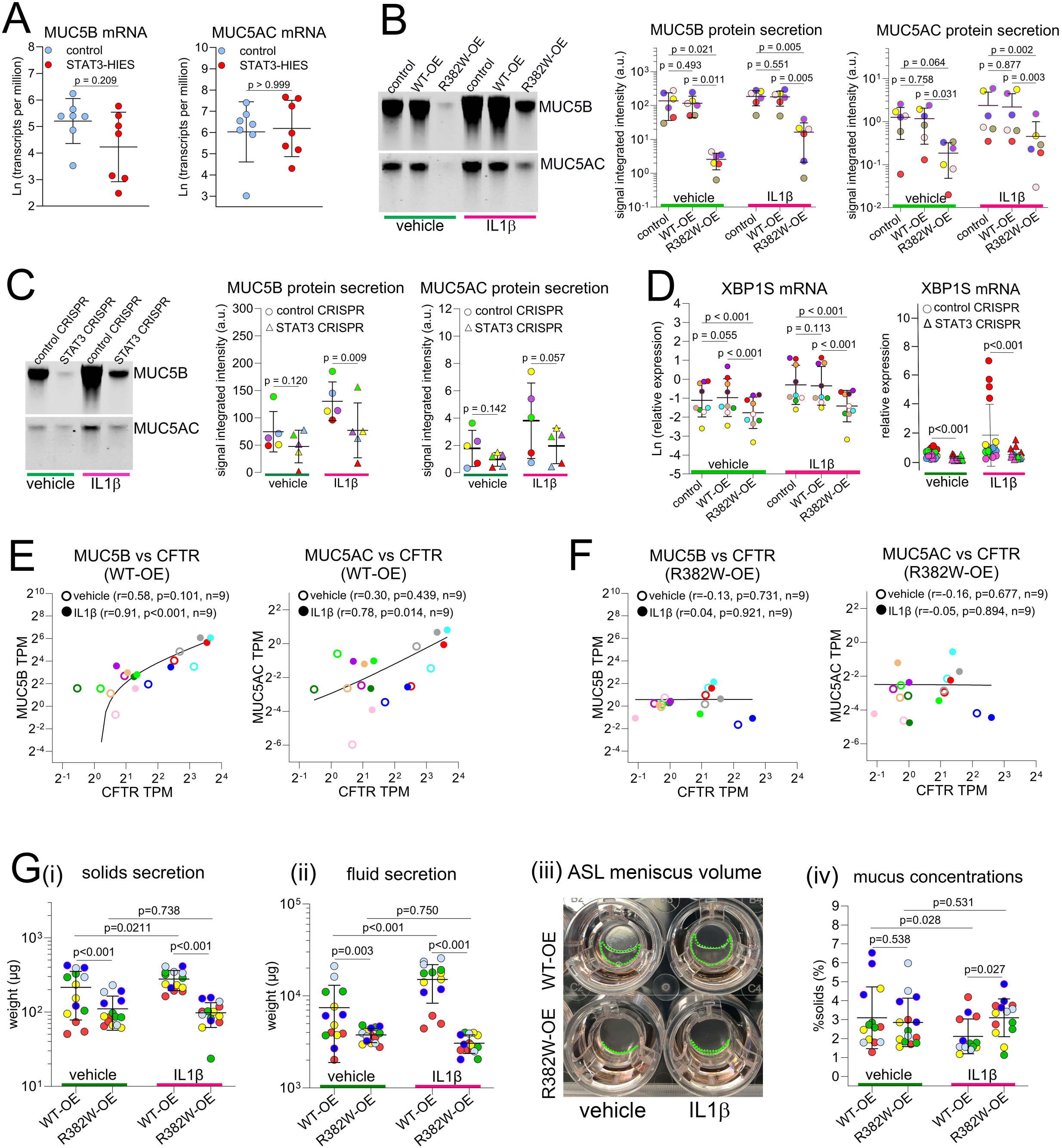
STAT3 is required for mucin and fluid secretions. **(A)** MUC5B and MUC5AC mRNAs of bronchial brushing tissue in vivo were quantified by bulk RNA-seq (7 donors/group). Normal HBE cells were transduced with **(B)** control, STAT3 WT-OE and STAT3 R382W-OE lentiviruses (n=6 donors), or **(C)** electroporated with control or STAT3 CRISPR/Cas9 (n=5 donors), followed by ALI culture. Apical secretion of MUC5B and MUC5AC proteins was detected by mucin agarose western blot and quantified by densitometry, and **(D)** the expression of XBP1S mRNA was determined by quantitative RT-PCR in the STAT3 R382W-OE and STAT3-KO HBE cells, respectively. **(E, F)** Linear regression tests were performed on CFTR mRNA with MUC5B and MUC5AC mRNAs (determined by bulk RNA-seq) in the normal HBE cells (n=9) transduced with **(E)** WT-OE vs **(F)** R382W-OE lentiviruses at baseline or after exposure to IL1β. **(G)** Apical **(i)** solids contents, (i.e., dry weight) and **(ii)** fluid (wet weight) secretions were measured in the control, WT-OE and R382W-OE HBE using microbalance. **(iii)** To visualize the effect of R382W-OE on apical fluid secretion, the culture plate was tilted to pool secretions at the Transwell’s bottom edge. The accumulated ASL meniscus volumes were highlighted by green dash lines in WT-OE vs R382W-OE HBE cells with or without IL1β treatment. **(iv)** Mucus concentrations (%solids) were calculated based on the ratio of dry to wet weight. Symbols with the same colors in the same panel indicate multiple cultures originated from the same donor, regardless of the shapes, except in **(A)**. HBE: human bronchial epithelia; ALI: air-liquid interface; Ln: natural logarithm; TPM: transcripts per million; ASL: airway surface liquid; OE: overexpression.

Consistent with STAT3 functions as a regulator of mucin and CFTR-mediated fluid secretion, R382W-OE exhibited decreased secretion of solids content (mucins, globular proteins, and salts) (Figure 4Gi) and fluid (Figure 4Gii,iii) compared to WT-OE HBE cells at baseline and post-IL1β. As a consequence of reduced secretion of both solids and fluid, R382W-OE HBE cells exhibited comparable ASL mucus concentration (%solids) at baseline. In response to IL1β, mucus concentration decreased in WT-OE, but not in R382W-OE HBE cells (Figure 4Giv), indicating that the relatively larger increase in fluid vs solids from WT-OE in response to IL1β was lost in R382W-OE HBE cells. These data suggest STAT3 deficiency compromises both fluid and mucin secretion, producing a unique condition of ASL volume depletion and modestly increased mucus concentrations in response to inflammation.

### STAT3 regulates the expression of defense molecules and ion exchangers governing ASL pH via IL1R1-dependent and independent pathways

Because STAT3-HIES lungs experience recurrent infection, the role of STAT3 mutations to regulate antimicrobial gene expression was investigated. The host defense gene lactotransferrin (LTF) (41) was decreased in STAT3-HIES bronchial brush specimens (Figure 5Ai). LTF mRNA was also decreased by STAT3 R382W-OE or KO in HBE cells at baseline and after IL1β exposure (Figure 5Aii,iii). STAT3 deficiency also decreased production of other biomolecules exhibiting antimicrobial activities under both basal and IL1β-stimulated conditions, including CCL20 (42), BPFIA1 (5, 43), LCN2 (44), and chemokines for recruitment of immune cells, CXCL1 and CSF3 (Figure 5B and Suppl Fig.7A,B). Further, STAT3 deficiency failed to induce a group of defense molecules after IL1β, but not at baseline, including IL17C (45) and AMP, e.g., β-defensin (DEFB4A) (Figure 5C, Suppl Fig. 7C), and components of the NADPH oxidase family (NOX1 and DUOX2) that produce microbiocidal reactive oxygen species (46) (Suppl Fig.7D). Notably, CRISPR/Cas9 ablation of IL1R1 also decreased the expression of these IL1β-regulated defense molecules in response to IL1β in primary HBE cells (Suppl Fig. 7E,F).

**Figure 5:**
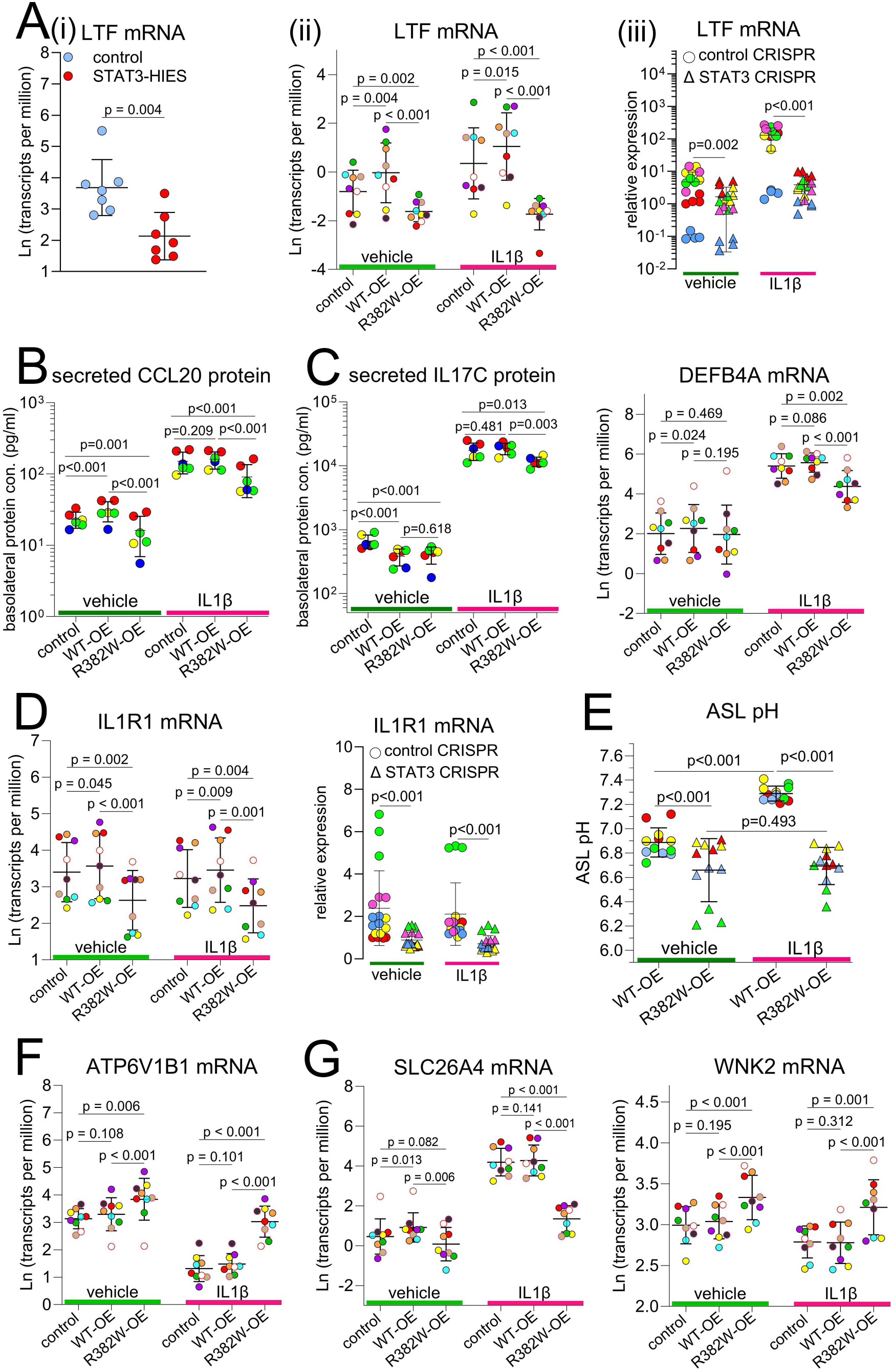
STAT3 regulates antimicrobial activities in IL1R1-dependent and -independent manners. **(A)** LTF (lactotransferrin) mRNA was quantified in **(i)** bronchial brushing specimens of control vs STAT3-HIES subjects by RNA-seq (7 donors/group). **(ii)** LTF mRNA expression in response to IL1β was measured in normal HBE cells (from 9 donors) expressing control, STAT3 WT-OE, and STAT3 R382W-OE lentiviruses, **(iii)** and STAT3-KO (by CRISPR/Cas9) HBE cells (from 5 donors) by bulk RNA-seq and TaqMan assays, respectively. **(B)** CCL20 (MIP-3α) and **(C)** IL17C secretion was measured by mesoscale U-PLEX assay platform in HBE cells (from 4 donors) expressing control, STAT3 WT-OE and STAT3 R382W-OE lentiviruses. **(C)** DEFB4A (β-defensin) mRNA expression in response to IL1β was measured in STAT3 R382W-OE HBE cells by bulk RNA-seq. **(D)** IL1R1 mRNA was measured in STAT3 R382W-OE and STAT3-KO HBE cells treated with or without IL1β by bulk RNA-seq and TaqMan assays, respectively. **(E)** ASL pH change in response to IL1β exposure was measured in the STAT3 WT-OE vs STAT3 R382W-OE HBE cells (from 4 donors) after 4 weeks of ALI culture. **(F)** mRNAs of proton transporter component ATP6V1B1, and **(G)** genes regulate bicarbonate secretions SLC26A4 and WNK2 in response to IL1β were measured by bulk RNA-seq in the HBE cells expressing control, STAT3 WT-OE and STAT3 R382W-OE lentiviruses. Symbols with the same colors in the same panel indicate multiple cultures originated from the same donor, regardless of the shapes. ASL: airway surface liquid; HBE: human bronchial epithelia; ALI: air-liquid interface; OE: overexpression.

Since STAT3 R382W-OE phenocopied IL1R1-KO regulation of defense genes, the role of STAT3 to regulate IL1R1 expression was tested. Both STAT3 R382W-OE and KO inhibited IL1R1 mRNA expression, and STAT3 and IL1R1 mRNAs were tightly, positively correlated in primary HBE cells (Figure 5D, Suppl Fig.7G), suggesting a regulatory role of STAT3 to IL1R1 expression. Thus, we propose a model in which STAT3 is required for host defense via both IL1R1-dependent and independent pathways to promote antimicrobial activities in airway epithelium (Suppl Fig.7H).

### STAT3 deficiency airway surface liquid (ASL) acidification

Antimicrobial peptides typically function optimally at neutral to alkaline pHs. Defective CFTR impairs HCO_3_^-^ secretion and causes ASL acidification (47), leading us to test whether STAT3 deficiency produced abnormal ASL pH. STAT3 R382W-OE HBE cells exhibited lower ASL pH at baseline and failed to alkalize ASL in response to IL1β (48) compared to WT-OE (Figure 5E). Abnormalities beyond CFTR dysfunction may additionally regulate ASL pH. For example, STAT3 R382W-OE increased mRNAs of the B1 subunit of V-ATPase ATP6V1B1 (Figure 5F), that is responsible for trans-apical membrane H^+^ transport (49). Reciprocally, R382W-OE decreased expression of inflammation-induced HCO_3_^-^ transporters, including pendrin (SLC26A4) (50) and the Na^+^-dependent, electrogenic HCO_3_^-^ cotransporter (SLC4A4) (Figure 5G, Suppl Fig.7I) (51). R382W-OE also increased WNK2, which negatively regulates HCO_3_^-^ secretion (52) (Figure 5G).

### STAT3 deficiency impairs ciliated cell differentiation in STAT3-HIES lungs

Ciliated cell morphologies were grossly normal, but sporadic regional losses of cilia were observed in STAT3-HIES bronchial brush biopsies (Figure 6A). Analyses of bronchial brush bulk RNA-seq data suggested impaired ciliated cell, but not deuterosomal cell differentiation, in STAT3-HIES airway epithelia (Figure 6B, Suppl Table 2), consistent with brush biopsy histology and previous reports (5). Expression of axonemal dynein genes associated with ciliogenesis and pathogenesis of PCD (DNAH5, CCDC40, DNAI1, HYDIN and DNAAF5) (53) were decreased in STAT3-HIES epithelia in vivo (Figure 6C, Suppl Fig.8A). Transcription factors required for multiciliogenesis, e.g., TP73, RFX2 and FOXJ1 (54), were also decreased in STAT3-HIES airway epithelium in vivo (Suppl Fig.8B).

**Figure 6:**
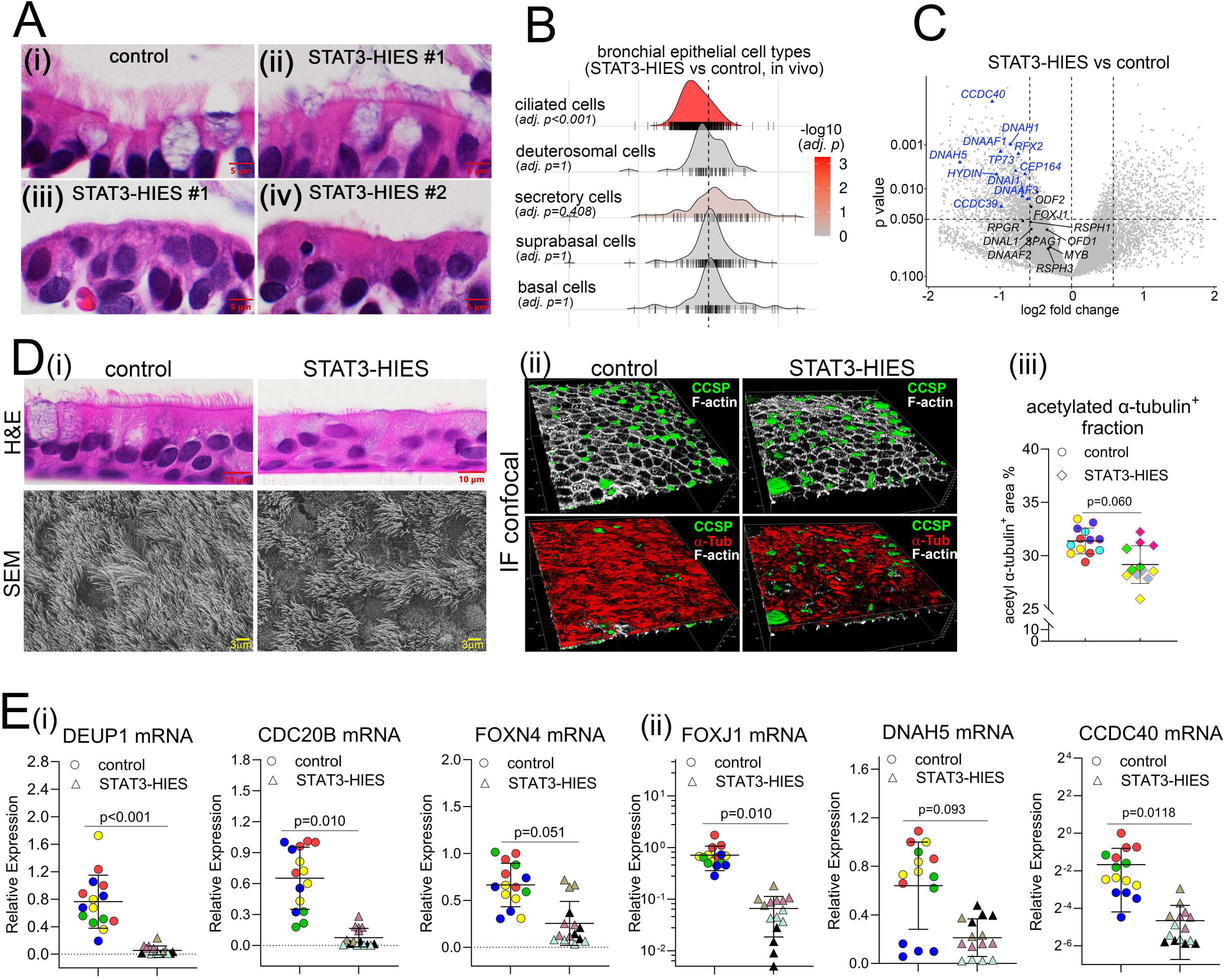
STAT3 mutations lead to impaired ciliogenesis in vivo and in vitro. **(A)** Representative histology of bronchial brushing tissues from **(i)** 1 control and 2 STAT3-HIES subjects [areas containing **(ii)** normal and **(iii, iv)** abnormal ciliated cell morphology] were shown by H&E staining. Images are representative of 3 donors/group. **(B)** Ridge plot represents the epithelial cell types from bulk RNA-seq of bronchial brushing tissues. Each ridge represents the enriched gene expression profile of one epithelial cell type, with red color intensity indicating adjusted p-values. The x-axis displays the log2 normalized expression fold change, and the y-axis displays the number of cells. Cell types were classified using the online published single cell RNA-seq database after enrichment analysis of gene ontology terms. **(C)** A Volcano plot displays differential gene expression analysis results from bulk RNA-seq of bronchial brushing tissues from STAT3-HIES vs control subjects (7 donors/group). A group of axonemal dynein genes related to ciliogenesis and PCD pathogenesis were analyzed for their expressions. Gene names listed in blue color indicate their changes are >-1.5 fold and p values <0.05. **(D) (i)** Epithelial cell histology of in vitro cultured control and STAT3-HIES HBE cells was shown by H&E staining (images were representative of 3 donors/group) and by scanning electron microscopy to compare ciliated cell differentiation (1 donor/group). **(ii)** Expression of SCGB1A1, acetylated α-tubulin (α-Tub), and Phalloidin (F-actin) was detected by whole mount IF staining, followed by 3-D confocal imaging and **(iii)** α-Tubulin expression quantification by morphometrics (from 4 donors/group). **(E) (i)** Deuterosomal cell makers CDC20B, DEUP1, FOXN4 and **(ii)** multiciliated cell markers FOXJ1, DNAH5, and CCDC40 mRNAs were quantified in the cultured control vs STAT3-HIES HBE cells by TaqMan assays. Symbols with the same color in the same panel indicate cultures originated from the same donor, regardless of shapes. IF: immunofluorescence; PCD: primary ciliary dyskinesia.

### Ciliation in cultured STAT3-HIES airway cells

To test a STAT3 role to regulate multiciliogenesis independent of inflammation/infection status in vivo, primary STAT3-HIES and control HBE cells were cultured under ALI conditions in vitro. STAT3-HIES cultures exhibited reduced ciliated cell numbers as demonstrated by histologic H&E staining, scanning electron microscopy, and α-tubulin immunohistochemistry (Figure 6D). Cultured STAT3-HIES HBE cells exhibited reduced deuterosomal and multiciliated cell gene expression (Figure 6E), suggesting deuterosomal cells might be regulated differently in vivo where inflammation stimulates their expansion (55).

### STAT3 R382W-mutation or KO HBE cells exhibit impaired ciliation

To test whether the JAK/STAT3 pathway is required for ciliogenesis, properties of STAT3 and JAK1 deficient cells were analyzed. STAT3 WT-OE HBE cells exhibited moderately increased % α-tubulin^+^ areas compared to controls (Figure 7A). In contrast, R382W-OE HBE cells exhibited reduced % α-tubulin^+^ areas compared to controls, consistent with reduced cilia formation at 4-week of ALI. RNA-seq demonstrated that at an early differentiation stage (ALI 8-day), STAT3 R382W-OE led to decreased expression of deuterosomal and multiciliated cell markers (Suppl Fig.9A). CRISPR/Cas9 ablation of STAT3 or JAK1 phenocopied R382W-OE by decreasing deuterosomal cell differentiation and mature ciliated cell numbers (Figure 7B, Suppl Fig.9B,C), suggesting the JAK1/STAT3 pathway is required for ciliogenesis.

**Figure 7:**
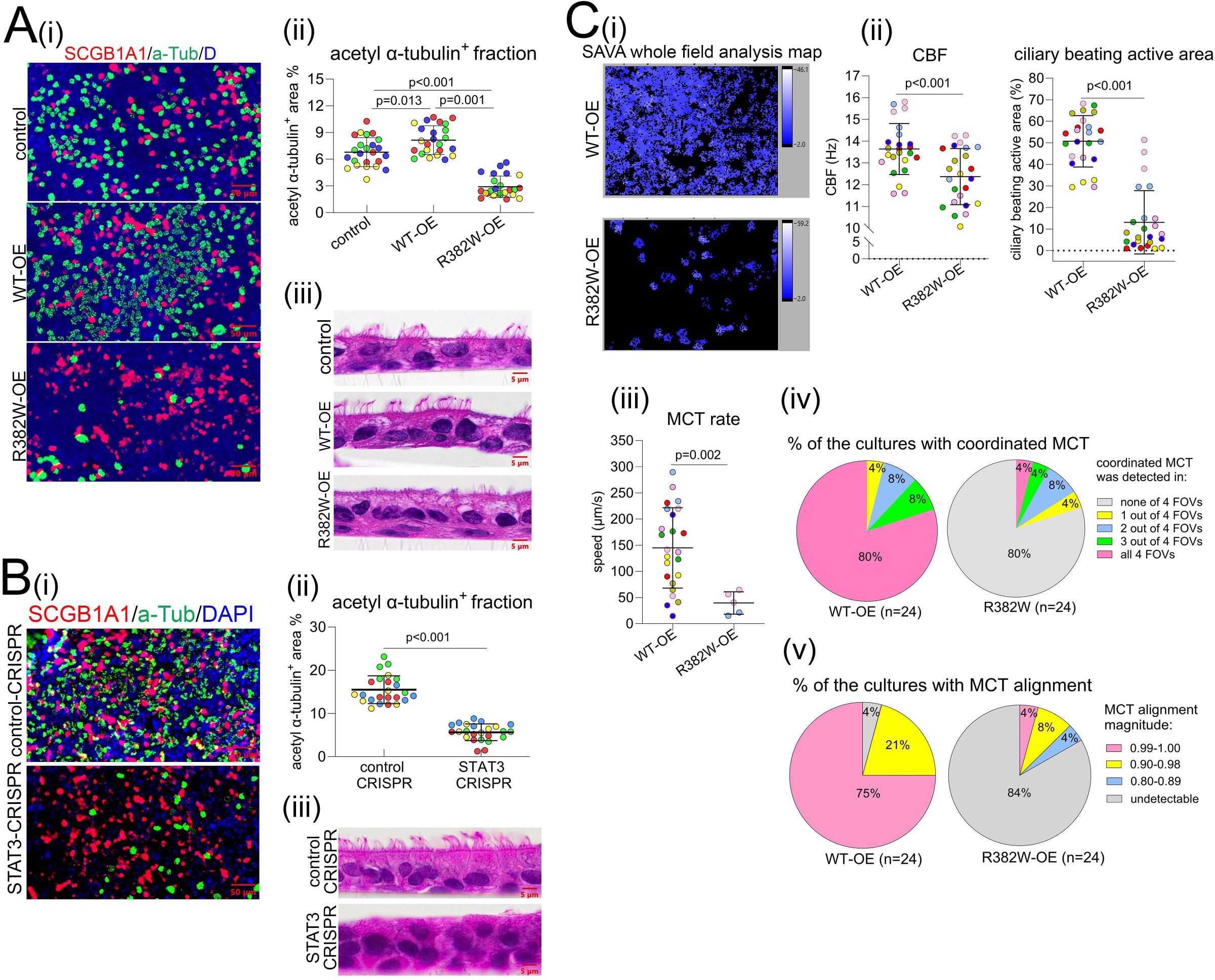
STAT3 R382W mutation or STAT3 KO leads to impaired ciliated cell differentiation and functions. **(A) (i)** Expression of SCGB1A1 and acetylated α-Tubulin (α-Tub) was detected by whole mount IF, followed by **(ii)** morphometric quantification of % area of α-Tubulin expression in the control, STAT3 WT-OE, and STAT3 R382W-OE lentiviruses infected HBE cultures (from 4 donors). **(iii)** Histology of these cells was shown by H&E staining. **(B)** Normal HBE cells (from 4 donors) were electroporated with control and STAT3 CRISPR/Cas9, **(i)** SCGB1A1 and α-Tub proteins were detected by whole mount IF followed by **(ii)** morphometric quantification. **(iii)** Histology of these cells was shown by H&E staining. **(C)** STAT3 WT-OE and STAT3 R382W-OE lentiviruses infected HBE cells were seeded on homemade racetrack to promote coordinated mucociliary transport. Ciliary beating activities were recorded by high-speed video microscopy and analyzed by Sisson-Ammons Video Analysis (SAVA) software. **(i)** The SAVA whole field view was shown to compare active ciliary beating between STAT3 WT-OE and R382W-OE cells. **(ii)** Ciliary beating frequency (CBF) and percentage of ciliary beating active area (surface ciliation) were measured and analyzed. **(iii)** Fluorescent dye labeled beads were added to the cultures to measure mucociliary transport rate by calculating speed of beads transport in coordinated fashion. **(iv)** The % of the cultures exhibited coordinated MCT in the 4-FOV of each culture was compared between 2 genotypes and shown in pie chart. **(v)** The alignment of the tracks of beads were measured and assigned to 4 magnitudes (fully aligned=1). The % of cultures that exhibited magnitude of alignment in each category were shown by pie charts. Symbols with the same color in the same panel indicate cultures originated from the same donor. IF: immunofluorescence; HBE: human bronchial epithelial; OE: overexpression; FOV: fields of view; MCT: mucociliary transport. All whole mount and histology images are representative of n≥3 donor cells.

### Cilia and MCC transport as a function of STAT3 function

To test whether defective STAT3 impairs MCC, cilial parameters and MCC were quantitated in STAT3 R382W-OE vs WT-OE HBE cells. R382W-OE exhibited reduced ciliation, modestly decreased CBF (~11%), lack of cilial beat coordination, alignment, and profoundly decreased vectorial MCC rates as compared to WT-OE HBE cells (Figure 7C, Suppl Fig.10 and Suppl Movie 1-5).

### Inhibition of the Notch pathway by γ-secretase inhibitor (GSI) restores ciliogenesis in STAT3 R382W-OE cells

Inhibition of Notch signaling is required for the progenitor to deuterosomal cell transition prior to multiciliogenesis (27, 56). STAT3 R382W-OE and WT-OE HBE cells were compared to identify genes known to regulate Notch activity during this transitional phase (Suppl Fig.11A). R382W-OE HBE cells exhibited reduced expression of HES6 (Figure 8A), a Notch pathway suppressor expressed during the transition from secretory cell progenitor to deuterosomal cell (27). Blocking Notch with GSIs promotes differentiation toward the multiciliated lineage, and GSI LY450139 (Semagacestat) increases multiciliogenesis in human nasal epithelium (57). To test whether LY450139 restored defective ciliogenesis in STAT3-HIES airway epithelium, R283W-OE HBE cells were used as a model for testing LY450139 efficacy at 3 to 5 weeks following ALI culture using concentrations reported to promote cilia structure and functions (57). LY450139 (both 250 and 500nM) increased HES6 and suppressed NOTCH1/3 (58) mRNA in R382W-OE cells (Figure 8B, Suppl Fig.11B), indicating Notch was inhibited. Importantly, expression of markers of deuterosomal and multiciliated cells, acetylated α-tubulin^+^ staining, ciliary active area, and CBF were increased in R382W-OE cells administered with LY450139 (Figure 8C-E). LY450139 administration did not decrease markers of secretory cell (CFTR or MUC5B), basal or suprabasal cell expression (Suppl Fig.11C,D). These data suggest LY450139 (250nM) improves multiciliogenesis without loss of secretory capacities in R382W-OE epithelia.

**Figure 8:**
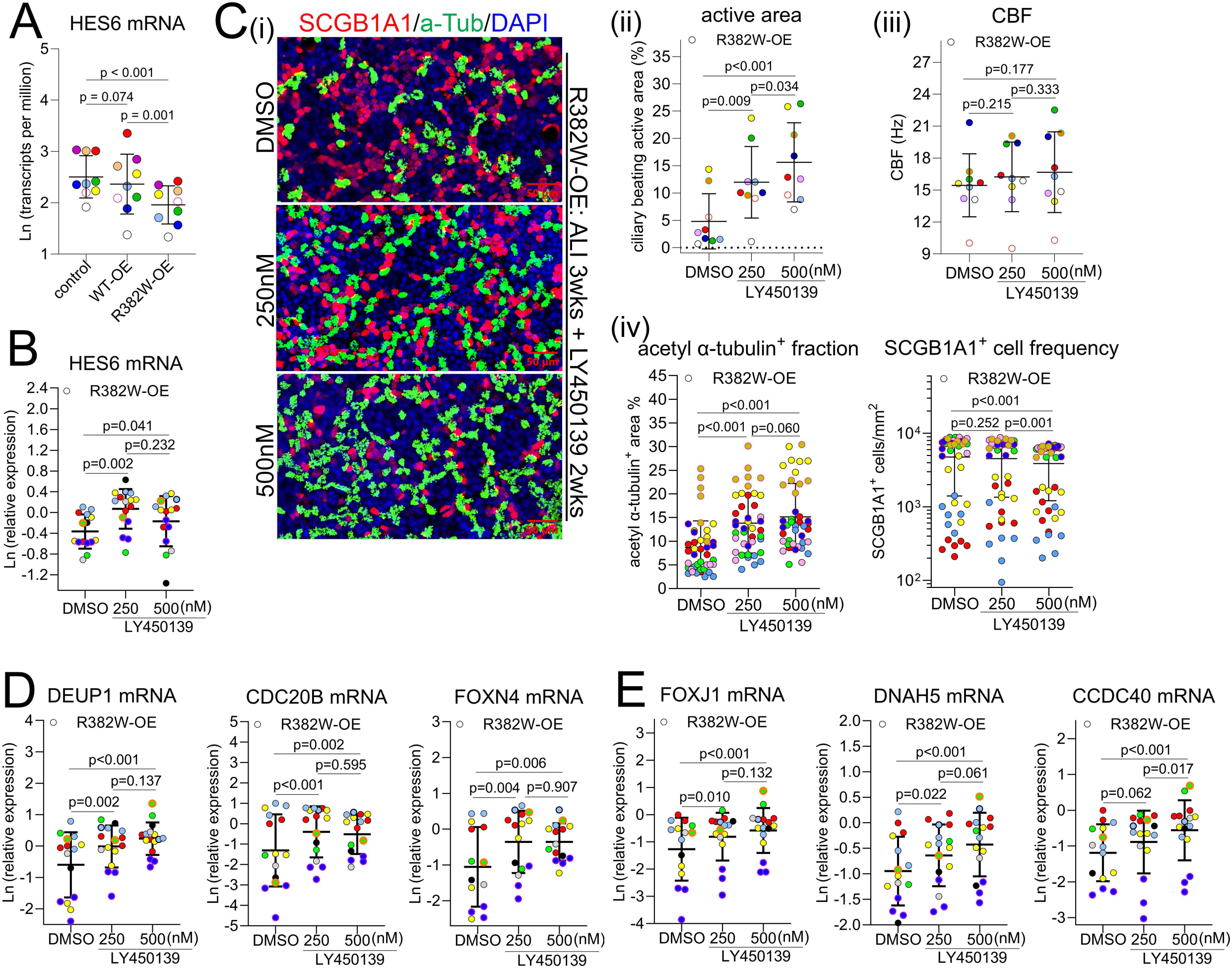
Gamma secretase inhibitor LY450139 increases HES6 expression and improves ciliogenesis in the STAT3 R382W mutant HBE cells. **(A)** HES6 mRNA was detected by bulk RNA-seq in the normal HBE cells (from 9 donors) expressing control, STAT3 WT-OE, and R382W-OE lentiviruses. **(B-E)** Normal HBE cells were transduced with STAT3 R382W-OE lentivirus and cultured under ALI conditions for 3 weeks prior to treatment with LY450139 (250, 500 nM) or DMSO for the following 2 weeks (n=9 donors). **(B)** Expression of HES6 mRNA in STAT3 R382W-OE HBE cells (from 7 donors) was determined by TaqMan assays. **(C) (i)** Expression of SCGB1A1 and acetylated α-tubulin (α-Tub) was detected by whole mount IF staining in STAT3 R382W-OE HBE cells (from 7 donors). **(ii)** Ciliary beating active area (surface ciliation) and **(iii)** CBF were measured by high-speed video microscopy in STAT3 R382W-OE HBE cells (from 9 donors). **(iv)** Acetylated α-Tub^+^ fraction and SCGB1A1+ cell frequency were determined by morphometrics of whole mount staining (n=7 donors). Expression of **(D)** the deuterosomal cell markers DEUP1, CDC20B, FOXN4 and **(E)** multiciliated cell makers FOXJ1, DNAH5, and CCDC40 mRNAs in the LY450139 treated STAT3 R382W-OE HBE cells (from 7 donors) was determined by TaqMan assays. Symbols with the same color in the same panel indicate cultures originated from the same donor HBE: human bronchial epithelial. IF: immunofluorescence; HBE: human bronchial epithelial; OE: overexpression; CBF: ciliary beating frequency.

## Discussion

Because of the well-described role of STAT3 to regulate immune cell differentiation and function (59), STAT3-HIES lung disease is generally considered to be immune cell mediated. However, our data and others (5) suggest that STAT3 dysfunction impairs not only immune cell function but also multiple aspects of airway epithelial host defense activities under baseline and injury/inflammatory conditions.

MCC is the dominant airway epithelial host defense mechanism against microorganisms (9), requiring the coordinated regulation of ASL hydration, pH, mucin secretion, and ciliary beat. STAT3-HIES airway epithelia exhibit defects in multiple components of the MCC apparatus. Like CF, STAT3-HIES airway epithelia exhibit impaired CFTR-dependent fluid secretion, but unique to STAT3-HIES, the change in CFTR expression is coupled with a reduction in mucin expression/secretion.

The consequences of reduced STAT3-HIES airway epithelial fluid and MUC5B secretion may parallel those observed in mice genetically altered to have decreased ASL volume (overexpression of β-ENaC transgene) and lack Muc5b expression (Muc5b^-/-^) (60). The β-ENaC/Muc5b^-/-^ mice exhibit worsened MCC, greater inflammation, and more severe bacterial infection than β-ENaC/Muc5b^+/+^ or WT mice (61). The worsening of MCC in the β-ENaC/Muc5b^-/-^ mice reflects low ASL volume coupled to the loss of the viscoelastic properties of Muc5b, while the worsened infection/inflammation may reflect that Muc5b is required to bind/organize secreted host defense proteins for optimal antimicrobial activity (62). Accordingly, the combination of reduced ASL volume and decreased MUC5B secretion in STAT3-HIES subjects may contribute to reduced MCC without dramatic increases in mucus concentration during the early disease phase.

The mucus property-dependent slowing of MCC in STAT3-HIES airways may be compounded by mutant STAT3-dependent defects in ciliated cell maturation. The degree of ciliation in fresh biopsies of STAT3-HIES subjects was variable, likely reflecting defects in regulation of transcriptional programs and signal pathways important for normal ciliogenesis and regeneration after injury (27). The STAT3 R382W-OE HBE cultures exhibited defects in ciliated cell maturation, as reflected in % of surface ciliation and modest CBF reductions, misaligned cilial orientation, and impaired vectorial MCC. Thus, STAT3-HIES airways may be vulnerable to collapse of MCC in response to injury, e.g., virus infections, that damage ciliated cells and produce limited ciliated cell repair (63, 64).

In coordination with the MCC apparatus, secretory cells secrete AMP (27) and recruit immune/inflammatory cells to suppress microbial proliferation until mechanical clearance by MCC is achieved (65). The data from STAT3-deficient HBEs suggest that STAT3-HIES subjects exhibit STAT3-dependent basal and IL1β-regulated defects in these activities, including: *1)* reduced mRNA expression of AMPs, e.g., lactotransferrin; *2)* reduced cytokine/chemokine protein secretion; and *3)* the inability to alkalize the ASL as required for apical surface antimicrobial activity.

A dominant regulator of airway secretory cell modification of mucus properties in disease is the IL1 cytokine family. Notably, each of the secretory cell activities in STAT3 deficient epithelia exhibited defective regulatory responses to IL1β as compared to STAT3 sufficient cells, including CFTR, mucin secretion, AMP, cytokines/chemokine signaling and ASL alkalization. The reduced expression of IL1R1 itself due to STAT3 dysfunction is an appealing unifying hypothesis for the failure of STAT3-HIES cells to mount appropriate host defense epithelial responses to IL1β.

The identification of a spectrum of innate host defense activities impaired in STAT3-HIES and STAT3 R283W-OE airway epithelia suggests a pathogenesis for the muco-obstructive, ultimately bronchiectatic, phenotype characteristic of STAT3-HIES subjects that emerges with age (66). Similar to CF (67), STAT3-HIES subjects do not exhibit lung disease at birth (68). However, as noted above (i.e., the ENaC/Muc5b^-/-^ model), STAT3-HIES lung may be more vulnerable to aspiration and infections during early-life due to deficient CFTR-mediated fluid and MUC5B secretion, which likely produces a low volume, low MUC5B content, poorly transportable mucus layer at birth. In parallel, the failure of AMP secretion and MUC5B-dependent AMP/host defense molecule organization on STAT3-HIES airway surfaces, with an acidic pH, may add to defective immune cells functions and favor robust bacterial growth in poorly cleared mucus. With time, repetitive aspiration and/or viral/bacterial-induced damage of cilia in STAT3-HIES airways impairs both the motor function of cilia required for mucus transport and the sensing function of cilia required to auto-regulate mucus concentrations. The failure to autoregulate mucus concentration promotes mucus hyperconcentration, and plugging, which initiates local epithelial hypoxic responses (69) that may further concentrate STAT3-HIES mucus into CF ranges. These defects generate vicious positive feedback cycles (see the proposed model in Figure 9) that ultimately produce bronchiectasis, and lung cysts, characteristic of late-stage STAT3-HIES (16).

**Figure 9:**
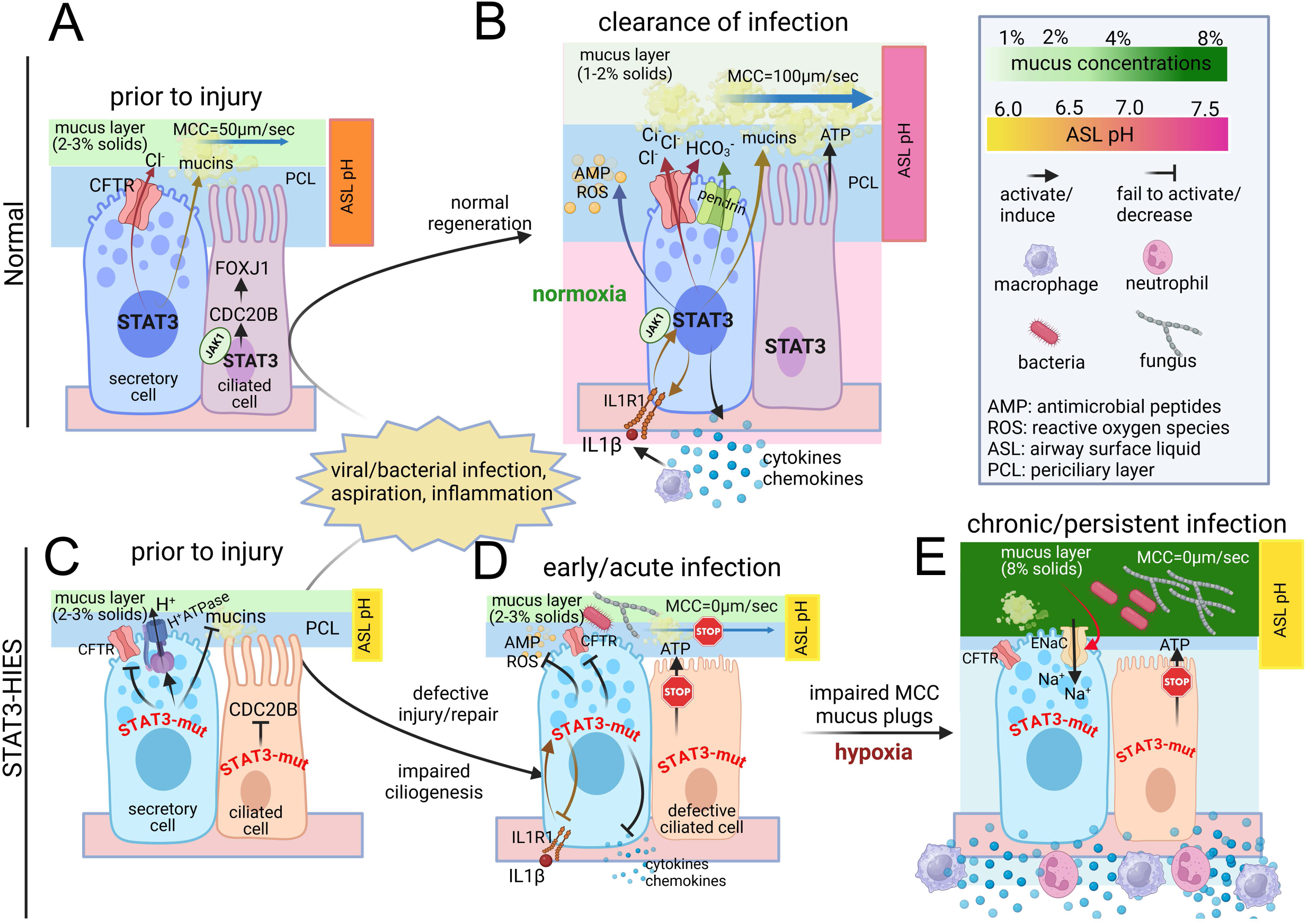
STAT3 deficiency leads to multiple-component defects in innate airway epithelial host defense, resulting in chronic infection. **(A)** In normal airway epithelial cells, STAT3 regulates basal CFTR-mediated fluid and mucin secretion, and JAK1/STAT3-mediated multiciliogenesis necessary for homeostatic mucociliary clearance (MCC) and normal airway surface liquid (ASL) pH. **(B)** In response to injuries (caused by infections, aspiration, and the resultant inflammation), STAT3 is activated by JAK1-mediated IL-1 signaling to increase CFTR-mediated fluid and mucin secretion, enhancing ASL volume while moderately decreasing mucus concentration compared to baseline. STAT3 is essential for IL1R1 expression, thus regulating the production and function of IL1R1-dependent and independent host defense molecules in response to IL1β. These include the secretion of antimicrobial peptides (AMP), reactive oxygen species (ROS), CFTR/pendrin-mediated bicarbonate (HCO3^-^) for ASL alkalization, and cytokines/chemokines critical for immune cell recruitment. These activities support coordinated MCC and microbicidal activities, clearing infections from the airways. **(C)** STAT3-HIES airway epithelial cells fail to maintain CFTR activity and mucin secretion, resulting in diminished ASL volume at baseline. Insufficient CFTR function, along with abnormal ion exchangers’ expression, leads to ASL acidification, increasing the vulnerability of STAT3-HIES epithelial cells. **(D)** Upon injuries, STAT3-HIES epithelial cells fail to adequately increase fluid and mucin secretion by secretory cells. The defective ciliated cell regeneration fails to provide motor function for MCC, resulting in low ASL volume, low MUC5B content, poor-transportable mucus layer, and static MCC. Decreased secretion of AMP/ROS, HCO3^-^ (for ASL alkalization), and cytokines/chemokines necessary for antimicrobial activity increase the susceptibility to infection on airway surface. **(E)** Impaired MCC during the acute phase leads to ongoing infection and inflammation, progressively increasing mucus concentration and leading to mucus hyperconcentration and plugging that produces local hypoxia in STAT3-HIES airways. Due to defective ciliation, the ATP-mediated autoregulation of mucus concentration via the mucus-cilia mechanism and motor function needed for mucus transport are lacking. Regional hypoxia promotes a vicious cycle by activating ENaC-mediated fluid/Na+ absorption and inhibiting ciliogenesis, that in turn further increasing mucus concentration and impairing MCC, resulting in chronic/persistent bacterial and fungal infection, and bronchiectasis over time.

The multiplicity of MCC defects in STAT3-HIES airways offers opportunities for novel therapies. GSI represents an option being explored therapeutically to block Notch and promote ciliogenesis (57), airway stem cell renewal, and epithelial barrier integrity in response to injury. LY450139 administration provides a therapeutic option for ciliated cell regeneration and restoration of MCC-mediated host defense in STAT3-HIES airways.

There are limitations to our study. First, STAT3-HIES is an extremely rare disease, and due to STAT3 mutant impairment of epithelial repair/proliferation (18), HBE cultures were generated from only four STAT3-HIES donors, which limited the statistical power of our findings. Second, the STAT3-genetically engineered R382W-OE HBE cultures used in our experiments do not fully capture the diversity of STAT3 heterozygous mutations present in STAT3-HIES subjects, nor do they replicate the chronic airway damage and inflammation status in vivo. Third, while our in vitro data suggest reduced ASL volume and mucin content in early-stage STAT3-HIES airways, these findings have yet to be validated with in vivo evidence. Finally, we tested host defense responses using IL1β as a surrogate for injury/inflammation stress. Although IL1β was elevated in STAT3-HIES BAL and highly induced in STAT3-HIES lungs, IL1β does not represent the entire inflammatory milieu in vivo that confronts abnormal STAT3-deficient epithelial cells.

In summary, STAT3 mediates a broad spectrum of host defense responses in human airway epithelial cells required for normal MCC and antimicrobial activities. The combined epithelial and immune cell host defense defects consequent to STAT3 mutations are consistent with the muco-obstructive, destructive lung disease phenotype that characterizes STAT3-HIES. Therapeutics directed at cilia defects of this syndrome, e.g., with Notch inhibitors, may be feasible.

## Acknowledgement

The authors thank Drs. Tom Ferkol (UNC at Chapel Hill) and Eszter Vladar (University of Colorado at Denver) for their insightful discussions that contributed to experiment design and manuscript writing, and Eric Roe for assistance with manuscript editing. They also thank Dr. Kathryn Ramsey (current affiliation: Telethon Kids Institute, Australia) for support in measuring mucus concentration in sputum, and Agathe Ceppe (UNC at Chapel Hill) for support with biostatistical analysis. Additionally, they acknowledge Leslie Fulcher and Allison Williams of the UNC Marsico Lung Institute Tissue Procurement and Cell Culture Core for providing normal human bronchial epithelial cells and reagents for cell culture, Susan Boyles of the CFTR Functional Core for CFTR activity measurements, and the UNC Center for Gastrointestinal Biology and Disease for histology services.

## Authors’ contributions

**LS** and **GC** designed and conceptualized, **LS**, **KNO**, **RCB** and **GC** supervised the project. **CR**, **AFF** and **KNO** provided patient care and collected clinical specimens from STAT3-HIES and health donors. **DH** and **AD** performed bioinformatic analysis and generated data. **XC** contributed to biostatistical analysis. **GR** and **MK** performed mucins mass spectrometry, **TGL** and **CRE** performed measurements of mucin sialic acid/urea concentrations, **MIG** and **BB** conducted mucus concentration measurements. **WJK**, **MRM** and **DBH** performed mucin viscoelasticity measurement in the sputum. **SHR** provided normal primary human bronchial epithelial (HBE) cells, advised on STAT3-HIES primary cell culture, experimental design, and manuscript writing. **LS** and **KH** performed measurements of mucus concentrations, bronchoalveolar lavage cytokine concentrations and airway surface liquid pH measurement. **SAW** performed Semagacestat study. **SAW**, **TS**, **FUD**, **MZ, DD**, **LZ** and **YM** performed immunofluorescent staining, imaging, coding for morphometric quantification. **LS** and **TS** performed treatments with pharmacological inhibitors, followed by quantification of proteins and mRNAs. **KO** performed RNA-seq of HBE cells, and **TA** performed IL1R1-KO by CRIPSR/Cas9 in HBE cells. **MC** performed 3D-confocal imaging; **CE** performed scanning electron microscopy to the cultured primary STAT3-HIES cells. **NLQ**, **TDR**, **MG** and **BRG** performed and analyzed CFTR function by Ussing chamber assays. **DMC** performed and quantified CFTR western blot. **SAW**, **PRS** and **LEO** analyzed ciliated cell differentiation and cilia functions. **LS** and **GC** drafted manuscript. **WKO**, **RB** and **GC** finalized the manuscript. All the authors read and approved the final manuscript.

## Supplementary Figure and Video Legends

**Supplementary Figure 1:**
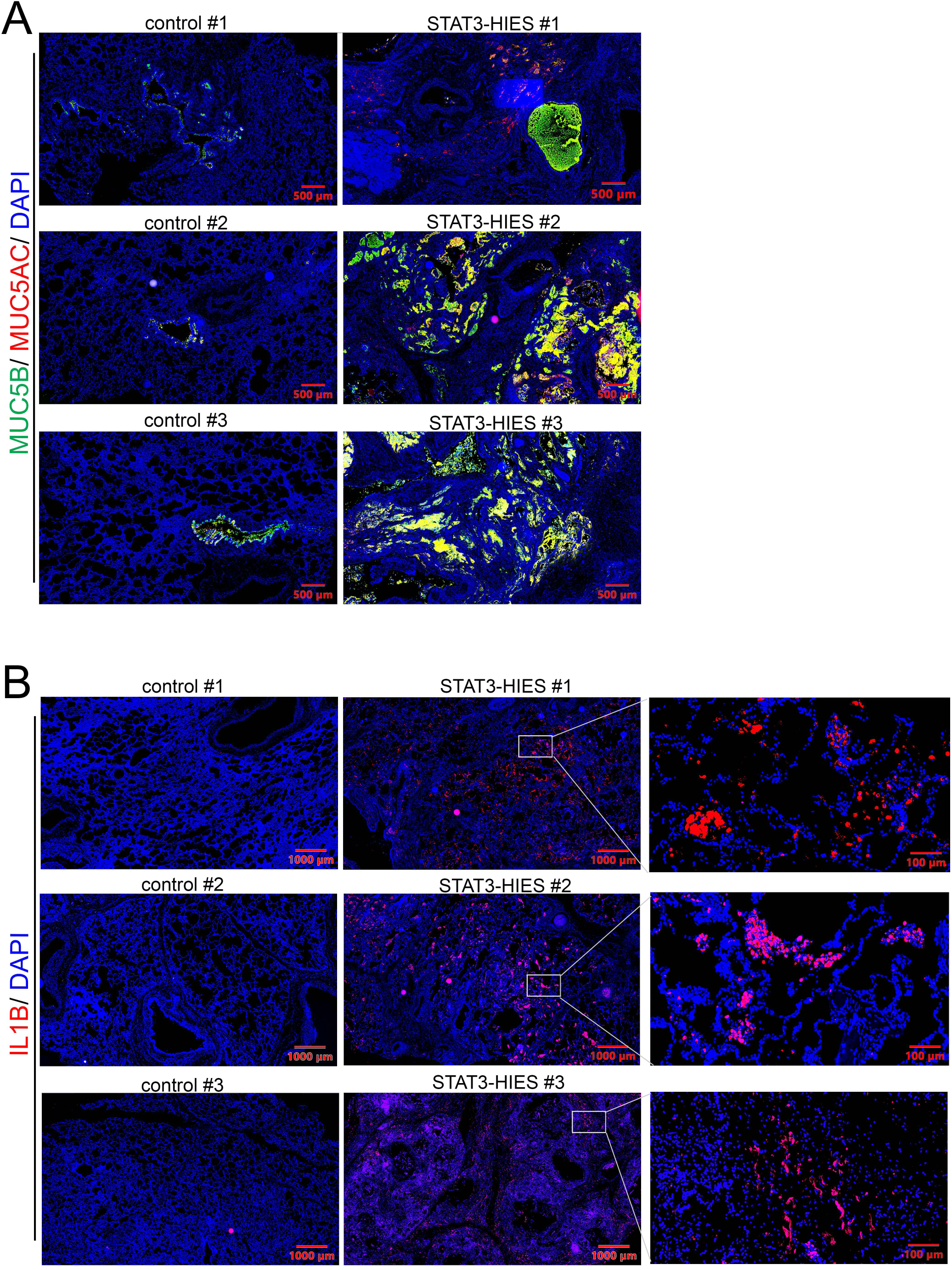
Expression of MUC5B, MUC5AC, and IL1B in control and STAT3-HIES lungs. **(A)** Expression of MUC5B (green) and MUC5AC (red) was detected by IF staining in the distal regions of the excised lung tissues from 3 control and 3 STAT3HIES donors. **(B)** Expression of IL1B (red) was detected by IF staining in the distal regions of the excised lung tissues from 3 control and 3 STAT3-HIES donors. The selected regions from STAT3-HIES lungs indicated by white rectangles are shown in higher power views on the right side of the low-power micrographs. IF: immunofluorescence.

**Supplementary Figure 2:**
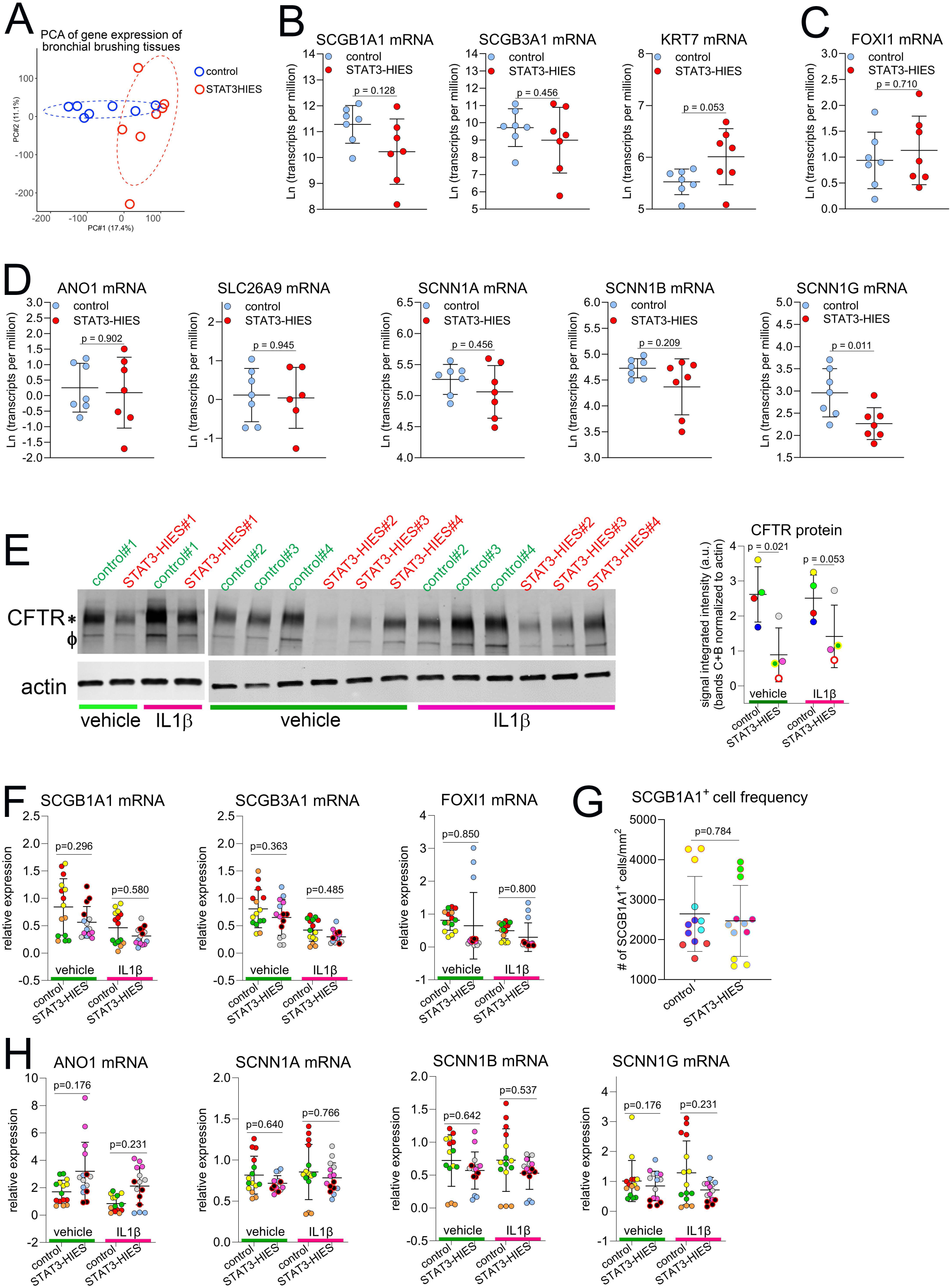
Expression of Club cell markers and ion transport channel genes in STAT3-HIES bronchial epithelial cells in vivo and in vitro. **(A)** Principal Component Analysis (PCA) plot of RNA-seq data showing the global gene expression profiles of bronchial brushing tissues from healthy controls vs STAT3-HIES donors (7 donors/group). Expression of mRNAs of **(B)** Club cell markers SCGB1A1, SCGB3A1, KRT7, **(C)** ionocyte marker FOXI1, and **(D)** epithelial ion channels ANO1 (TMEM16A), SLC26A9, SCNN1A, B, and G were quantified by bulk RNA-seq in control vs STAT3-HIES bronchial brushing tissues. **(E)** CFTR protein from primary control and STAT3-HIES HBE cells (4 donors/group) was measured by western blot followed by densitometry quantification. * and Φ represent band C and B of CFTR protein, respectively. Actin expression served as loading control. **(F-H)** Control and STAT3-HIES HBE cells (4 donors/group) were differentiated under ALI conditions prior to exposure with IL1β. **(F)** Expressions of SCGB1A1, SCGB3A1and FOXI1 mRNAs were determined by TaqMan assays and **(G)** the SCGB1A1^+^ cell frequencies presented in cultures were quantified by whole mount IF staining and morphometrics. **(H)** Expression of ion channel genes ANO1 (TMEM16A), SCNN1A, B, and G mRNAs was determined by TaqMan assays. Symbols with the same colors in the same panel indicate multiple cultures originated from the same donor, except in **(A-D)**. ALI: air-liquid interface; HBE: human bronchial epithelial: IF: immunofluorescence.

**Supplementary Figure 3:**
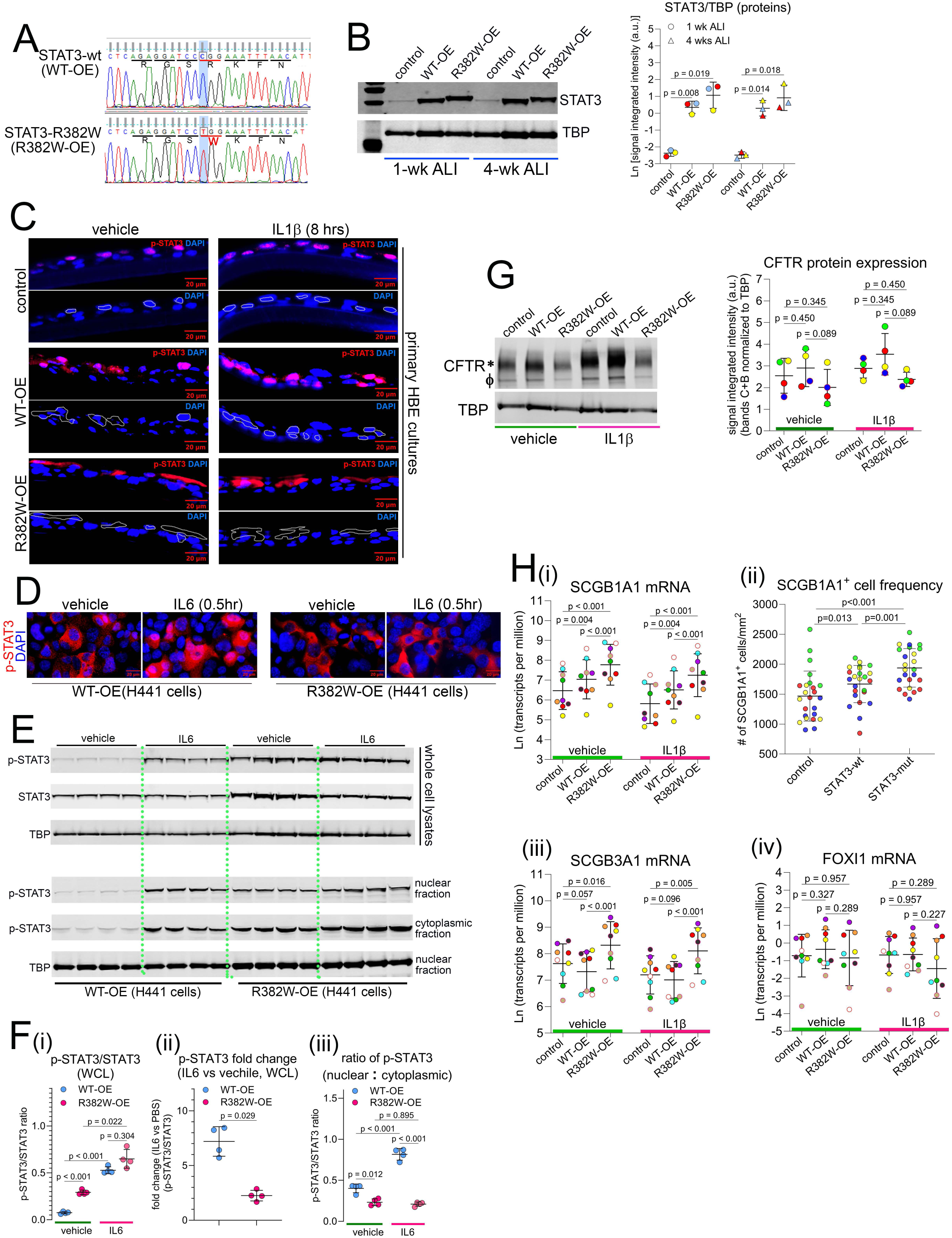
STAT3 R382W mutation reduces STAT3 phosphorylation efficiency in response to IL6, inhibits p-STAT3 nuclear translocation and decreases CFTR protein expression. **(A)** cDNA sequences of wild type STAT3 (STAT3 WT-OE) and STAT3 R382W mutation were cloned to pLVX-EF1α-IRES-Puro lentiviral vector. The R283W mutation was confirmed by Sanger sequencing. **(B)** Normal HBE cells (n=3 donors) transduced with control, STAT3 WT-OE and R382W-OE lentiviruses were cultured under ALI conditions. STAT3 protein expression at 1 and 4 weeks of ALI were detected by western blot and quantified by densitometry. The TATA box binding protein (TBP) served as a loading control. **(C)** Normal HBE cells (n=3 donors) were transduced with control, WT-OE and R382W-OE lentiviruses. The subcellular localization of p-STAT3 (upper panel of each section) in response to IL1β was detected by IF staining. The white dash-lines in the lower panels provided a visual guide for the spatial distribution of p-STAT3 within the subcellular architecture. **(D-E)** STAT3 WT-OE and R382W-OE lentiviruses were transduced into H441 cells. **(D)** The subcellular localization of p-STAT3 in H441 was detected by IF staining, and **(E)** expression of p-STAT3, STAT3 and TBP was detected by western blot in the whole cell lysates and in nuclear vs cytoplasmic fractions. **(F)** The western blot signals were semi-quantified by densitometry for calculation of **(i)** the ratio of p-STAT3/STAT3 in whole cell lysates (WCL), **(ii)** the fold change of p-STAT3/STAT3 in response to IL6 and **(iii)** the ratio of p-STAT3 in nuclear vs cytoplasmic fraction in response to IL6 in H441 cells. Experiments were repeated in 4 independent cultures. **(G)** CFTR protein in control, WT-OE and R382W-OE lentiviruses infected HBE cultures was detected by western blot. The image is representative of HBE cells from 4 donors. The signals on the western blot were semi-quantified by densitometry. **(Hi,iii,iv)** SCGB1A1, SCGB3A1 and FOXI1 mRNAs were quantified by RNA-seq. **(Hii)** The frequency of SCGB1A1^+^ cells was quantified by whole mount staining and morphometrics. Symbols with the same colors in the same panel indicate multiple cultures originated from the same donor. IF: immunofluorescence; ALI: air-liquid interface; HBE: human bronchial epithelial; WCL: whole cell lysates, OE: overexpression.

**Supplementary Figure 4:**
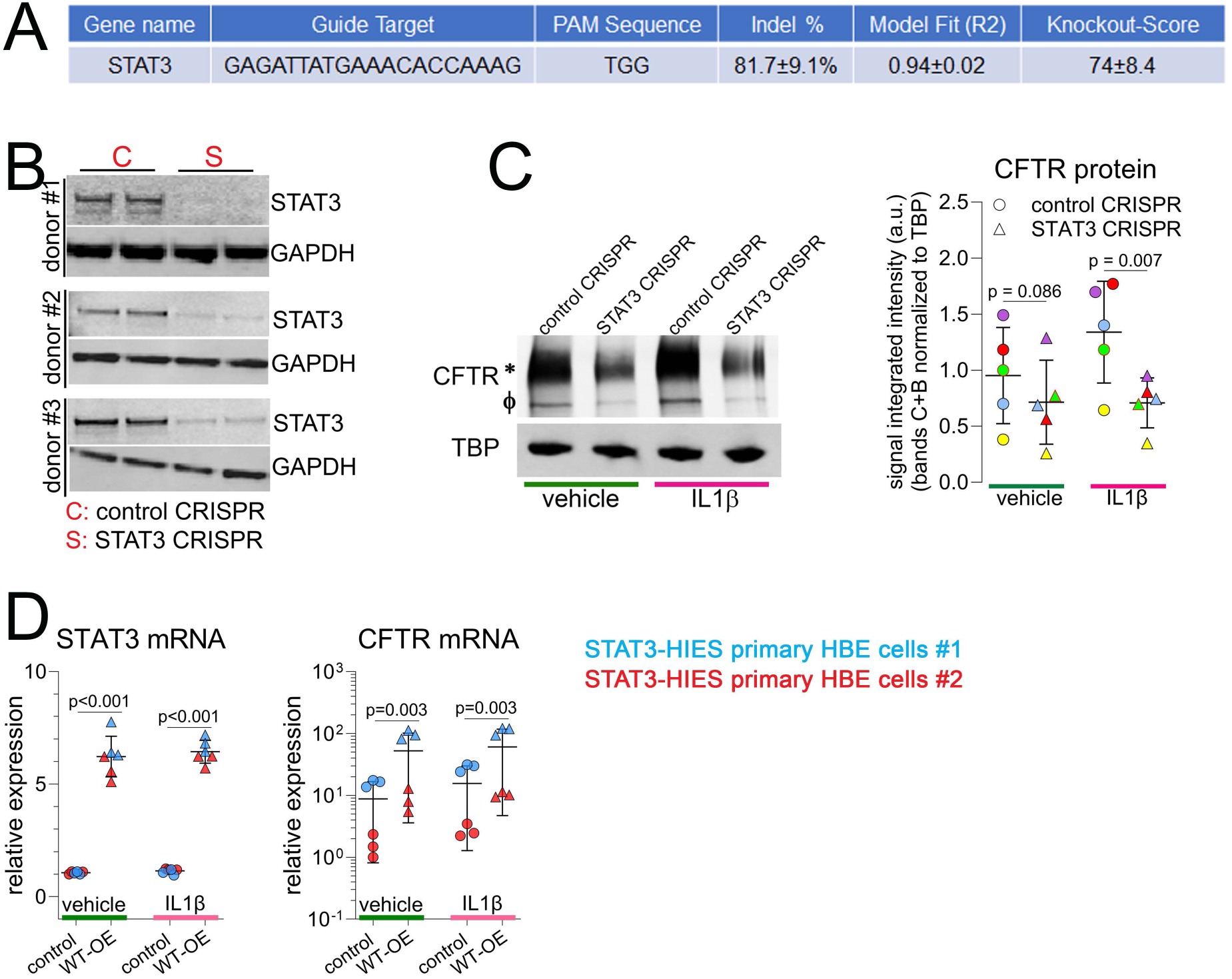
STAT3 knockout (KO) leads to reduced CFTR expression in normal HBE cells. **(A)** STAT3 CRISPR/Cas9 ribonucleoprotein complex was electroporated into HBE cells. Genomic DNA was isolated for Sanger sequencing and analyzed by Synthego online analysis tool. Expression of **(B)** STAT3 and **(C)** CFTR protein in control vs STAT3 KO HBE was detected by western blot. The images are representatives of western blots of HBE cells from 5 donors. **(D)** Primary STAT3-HIES HBE cells (from 2 donors) were transduced with control or STAT3 WT-OE lentiviruses followed by ALI culture. Expression of STAT3 and CFTR mRNAs was determined by TaqMan assays. Symbols with the same colors in the same panel indicate multiple cultures originated from the same donor regardless of shapes. KO: knockout; HBE: human bronchial epithelial; ALI: air-liquid interface; OE: overexpression.

**Supplementary Figure 5:**
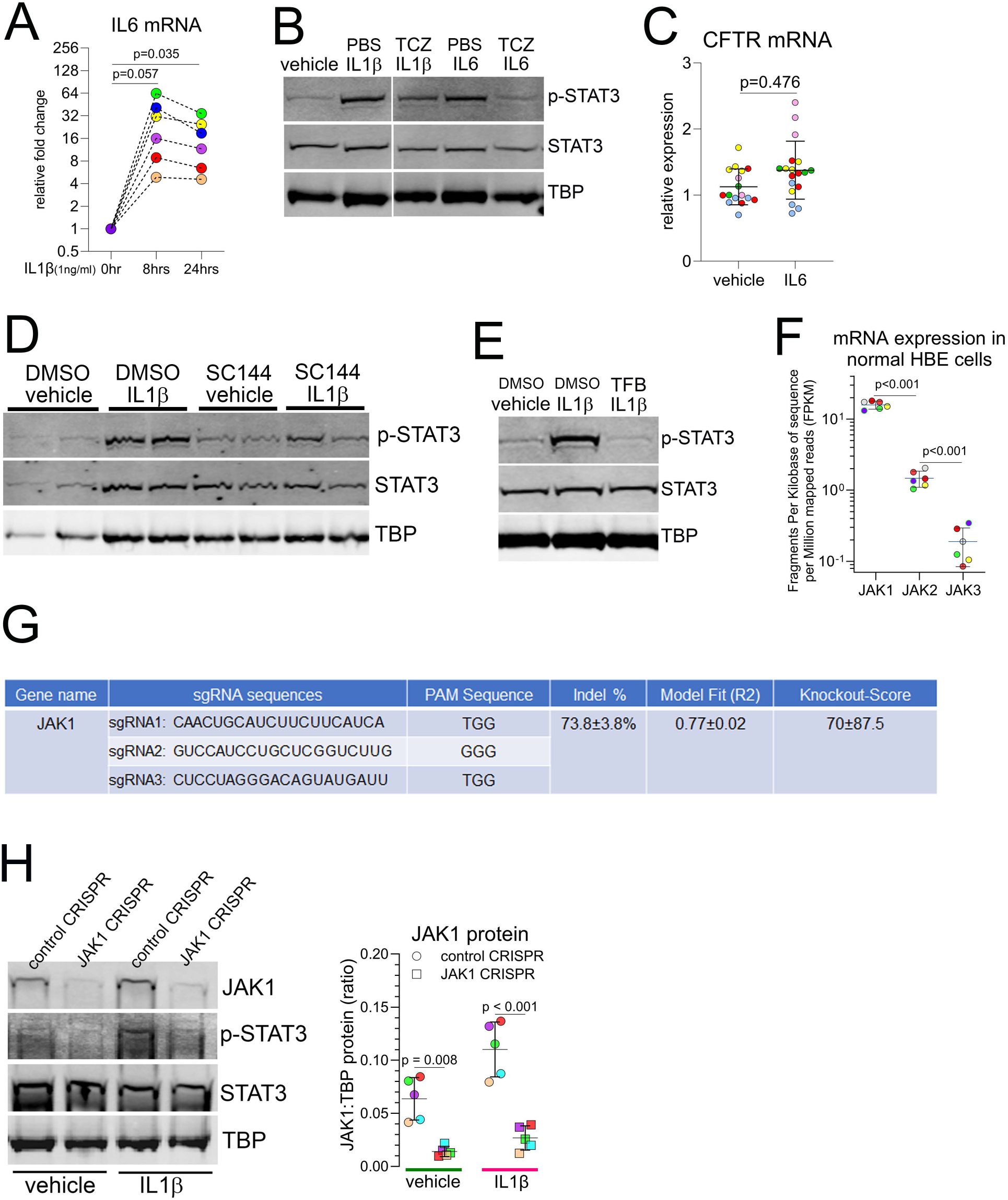
GP130-JAK1 signaling mediates IL1β-induced STAT3 phosphorylation and CFTR expression. **(A)** Normal HBE cells (n=6 donors) were exposed to IL1β for 8 and 24 hours, and IL6 mRNA fold change was compared to 0 hour by TaqMan assays. **(B)** Normal HBE cells were treated with or without IL6R monoclonal antibody Tocilizumab (TCZ) prior to exposure to IL1β or IL6 for 8 hours. p-STAT3 and STAT3 proteins were detected by western blot. The image is representative HBE cells from 8 donors. **(C)** CFTR mRNA in response to IL6 exposure was measured by TaqMan assays in well differentiated normal HBE cultures (n=5 donors). **(D)** Expression of p-STAT3 and STAT3 proteins was detected by western blot in normal HBE cells treated with GP130 inhibitor SC144 (10µM) or DMSO (vehicle) prior to expose to IL1β (1ng/ml) for 8 hours. The image is representative of HBE cells from 3 donors with 2 cultures/condition/donor. **(E)** p-STAT3, STAT3 proteins were detected by western blot in normal HBE cells pre-treated with JAK inhibitor Tofacitinib (TFB) prior to IL1β exposure. The image is a representative of HBE cells from 8 donors. **(F)** JAK1,2,3 mRNAs in normal HBE cells (n=6 donors) were determined by bulk RNA-seq and. **(G)** JAK1 CRISPRs were composed of combination of 3 individual CRISPRs shown in the table, and Indel%, Knockout-Score were calculated in targeted normal HBE. **(H)** Expression of JAK1, p-STAT3, STAT3 and TBP proteins was determined by western blot in HBE cells targeted by control or JAK1 CRISPR/Cas9 exposed to vehicle or IL1β. JAK1 expression was determined by western blot and semi-quantified by densitometry. The image is representative of HBE cells from 5 donors. Symbols with the same colors in the same panel indicate multiple cultures originated from the same donor. KO: knockout; ALI: air-liquid interface; HBE: human bronchial epithelial.

**Supplementary Figure 6:**
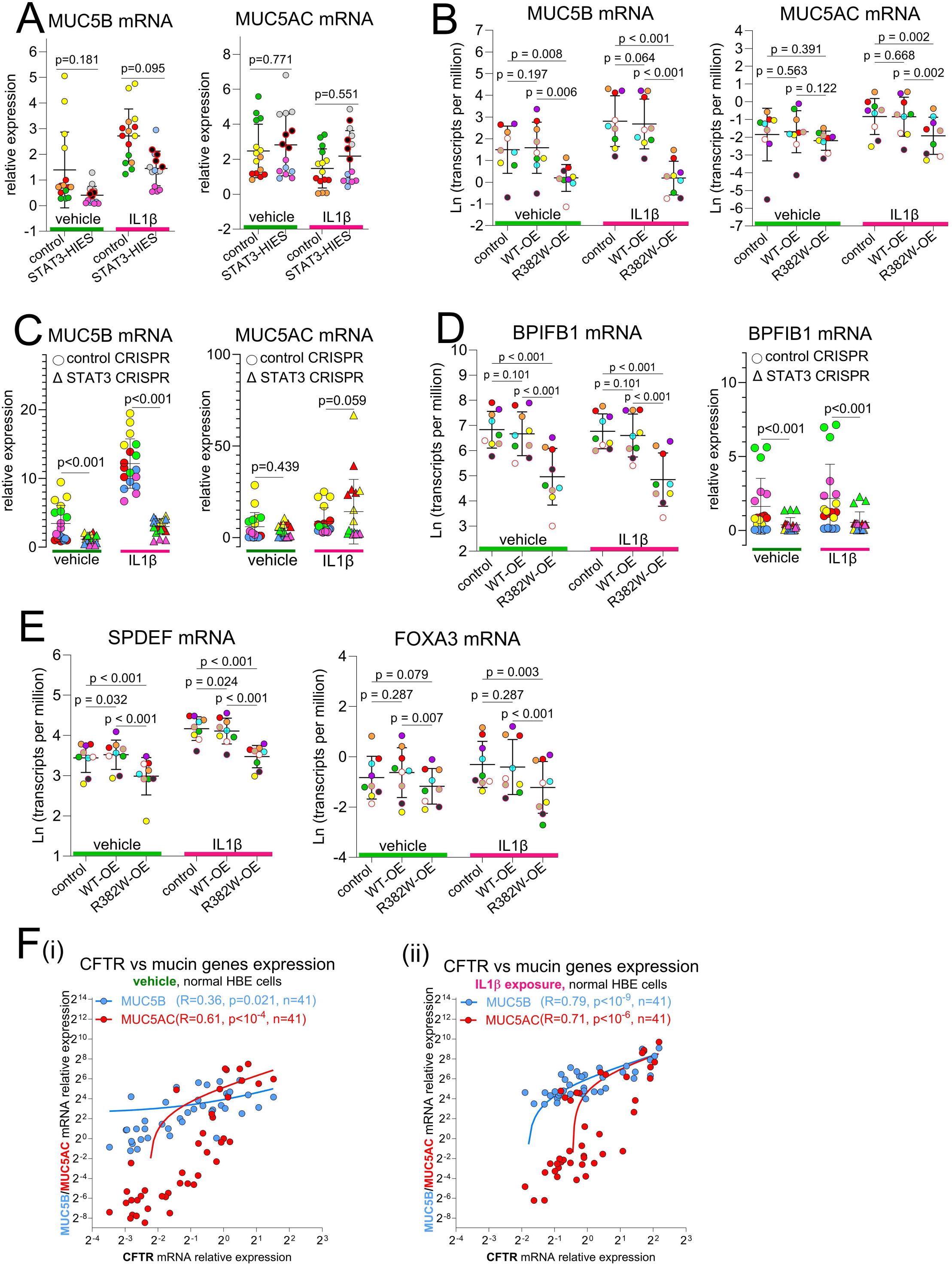
STAT3 deficiency leads to disproportional mucin vs CFTR expression in response to IL1β exposure. **(A)** MUC5B and MUCAC mRNAs were measured by TaqMan assays in control vs STAT3-HIES primary HBE cells (n=4 donors/group) cultured in vitro with or without IL1β exposure. **(B)** MUC5B and MUC5AC mRNAs were measured by bulk RNA-seq in the normal HBE cells (n=9 donors) expressing control, STAT3 WT-OE and STAT3 R382W-OE lentiviruses under ALI conditions prior to IL1β exposure. **(C)** MUC5B and MUC5AC mRNAs were measured in STAT3-KO (by CRISPR/Cas9) HBE (n=5 donors) cultured under ALI conditions prior to IL1β exposure. **(D)** BPIFB1 mRNA was measured in R382W-OE or STAT3 KO HBE. **(E)** SPDEF and FOXA3 mRNAs were quantified by bulk RNA-seq in normal HBE cells expressing control, WT-OE and R382W-OE lentiviruses, and treated with or without IL1β. **(F)** Basal and IL1β-induced CFTR, MUC5B, and MUC5AC mRNAs were quantified by TaqMan assays in normal well differentiated HBE from 41 donors. The correlations between CFTR (x-axis) and MUC5B or MUC5AC (shown in blue or red, respectively, both on the y-axis) mRNAs in normal HBE at **(i)** baseline and **(ii)** exposure to IL1β were demonstrated by linear regression. Symbols with the same colors in the same panel indicate multiple cultures originated from the same donor, regardless of the shapes, except in **(F)**. HBE: human bronchial epithelia; ALI: air-liquid interface; OE: overexpression.

**Supplementary Figure 7:**
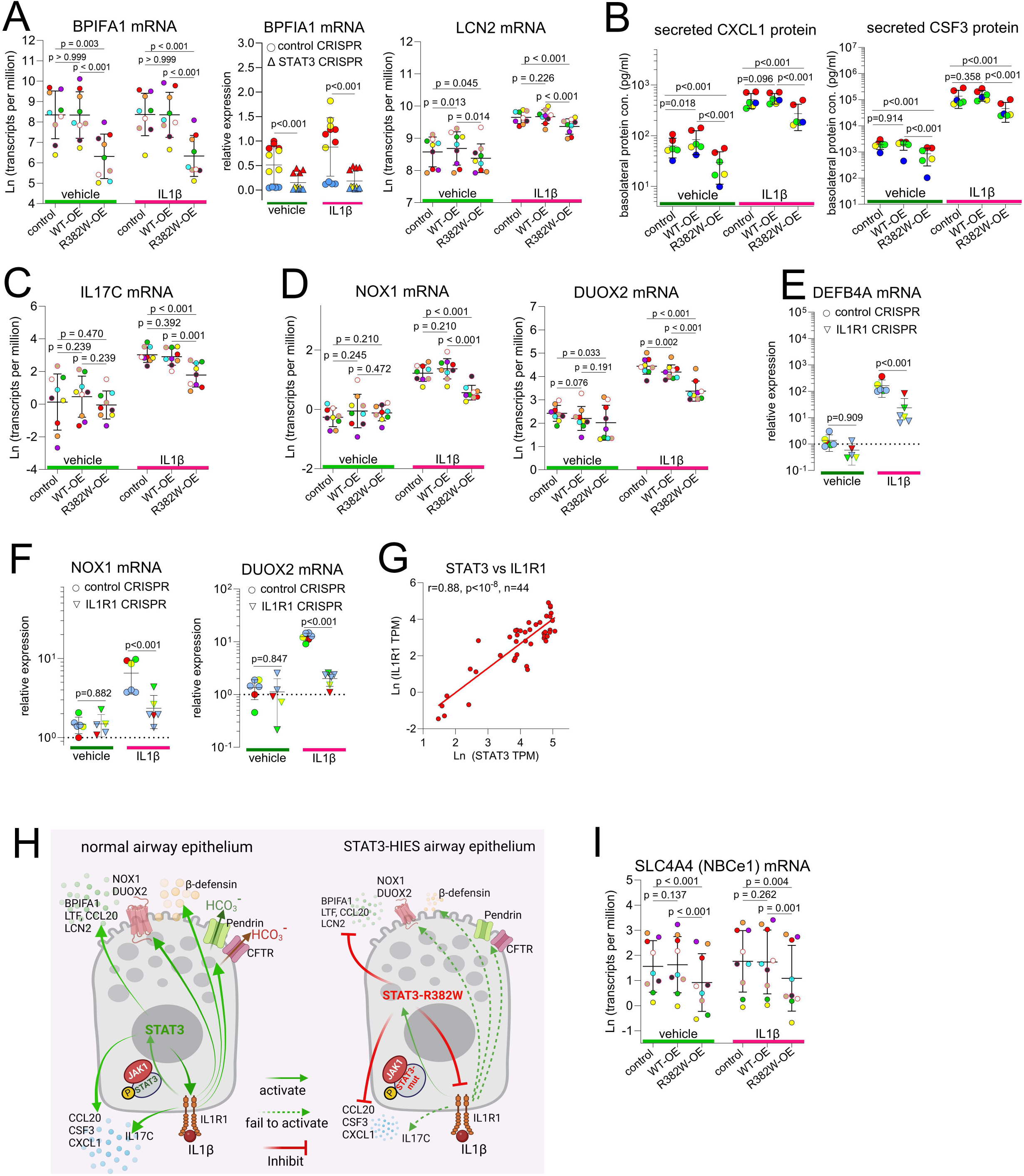
STAT3 regulates airway epithelial host defense in IL1R1-dependent and -independent manners. Normal HBE cells (from 9 donors) were transduced with control, STAT3 WT-OE, and STAT3 R382W-OE lentiviruses and cultured under ALI conditions prior to IL1β exposure. STAT3-KO HBE cells were generated by CRISPR/Cas9 and ALI cultured prior to IL1β exposure. **(A)** BPIFA1 mRNA was measured in R382W-OE and STAT3-KO, and their control HBE cells by RNA-seq and TaqMan assays, respectively. **(B)** Basolateral secretions of CXCL1 and CSF3 proteins were measured by mesoscale multiplex. **(C)** IL17C, **(D)** S100A8, S100A9, NOX1, and DUOX2 mRNAs were measured by bulk RNA-seq in the R382W-OE vs WT-OE and control HBE cells. **(E)** DEFB4A, **(F)** S100A8, S100A9, NOX1, and DUOX2 mRNAs were measured in IL1R1-KO (targeted by IL1R1 CRISPR/Cas9) HBE cells (from 4 donors). **(G)** The correlations between STAT3 and IL1R1 mRNAs (measured by bulk RNA-seq) in normal well differentiated HBE cells (n=44) were tested by linear regression. **(H)** A proposed model of STAT3 regulation of innate epithelial defense through IL1R1-dependent and -independent pathways: At baseline, STAT3 is required for producing antimicrobial molecules BPIFA1, LTF, CCL20, LCN2 and chemokines CXCL1, and CSF3. STAT3 is required for IL1R1 expression in HBE cells. Therefore, STAT3 controls expression of IL1β-inducible host defense molecules, including β-defensin, NOX1, DUOX2, IL17C, and genes regulate ASL pH, including CFTR, pendrin and other ion transporters. **(I)** Na^+^-dependent, HCO_3_^-^ electrogenic cotransporter (NBCe1), SLC4A4 mRNA was detected by bulk RNA-seq in normal HBE cells (n=9 donors) transduced with control, WT-OE, and R382W-OE lentiviruses exposed with or without IL1β. ASL: airway surface liquid; HBE: human bronchial epithelial; KO: knock-out; OE: overexpression; Ln: natural logarithm; TMP: transcripts per million.

**Supplementary Figure 8:**
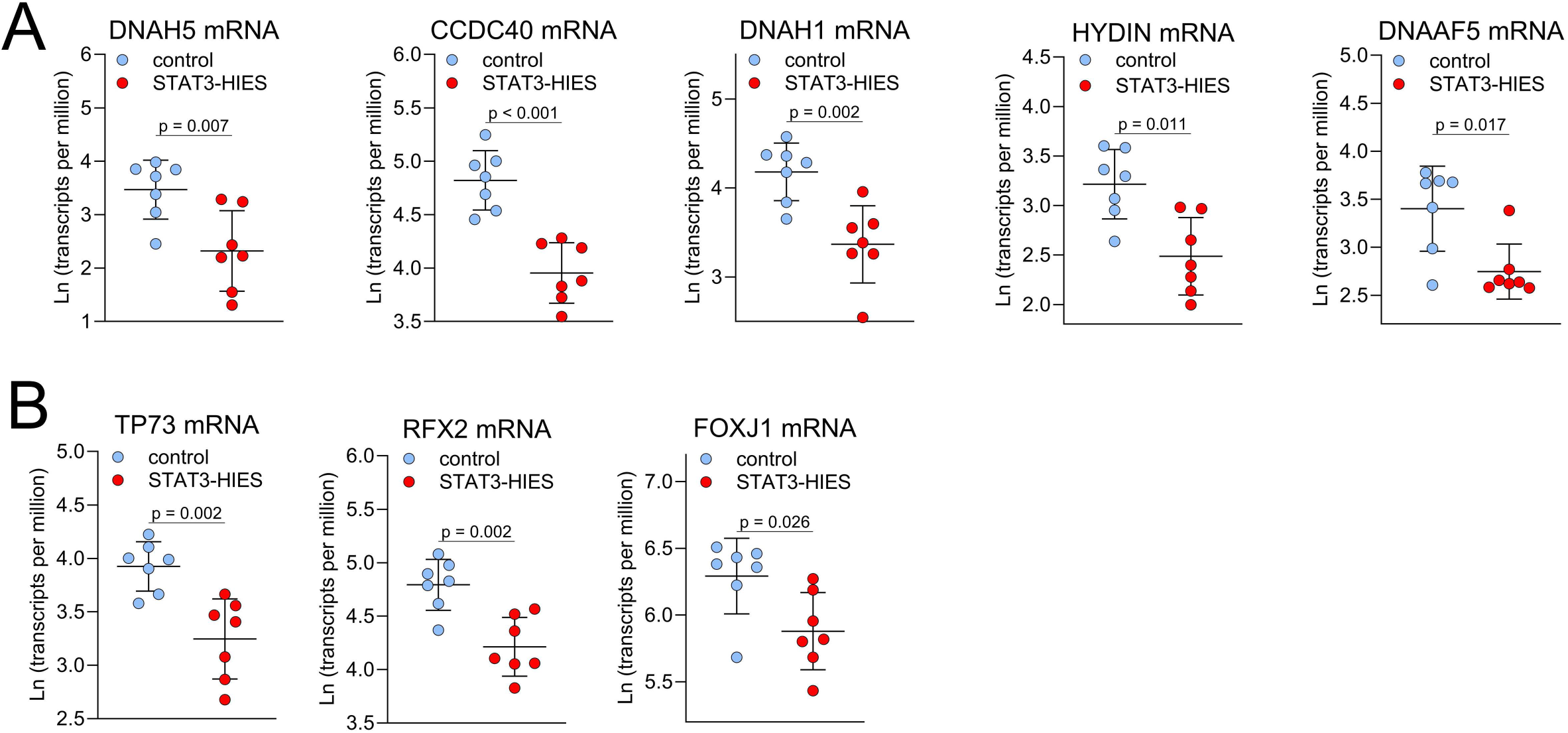
STAT3 deficiency leads to impaired differentiation of multiciliated cells in vivo and in vitro. **(A)** Dynein genes related to ciliogenesis and pathogenesis of PCD DNAH5, CCDC40, DNAH1, HYDIN, and DNAAF5 mRNAs, and **(B)** transcription factors TP73, RFX2, and FOXJ1 mRNAs were detected by bulk RNA-seq in bronchial brushing tissues of control vs STAT3-HIES donors (7 donors/group). PCD: primary ciliary dyskinesia.

**Supplementary Figure 9:**
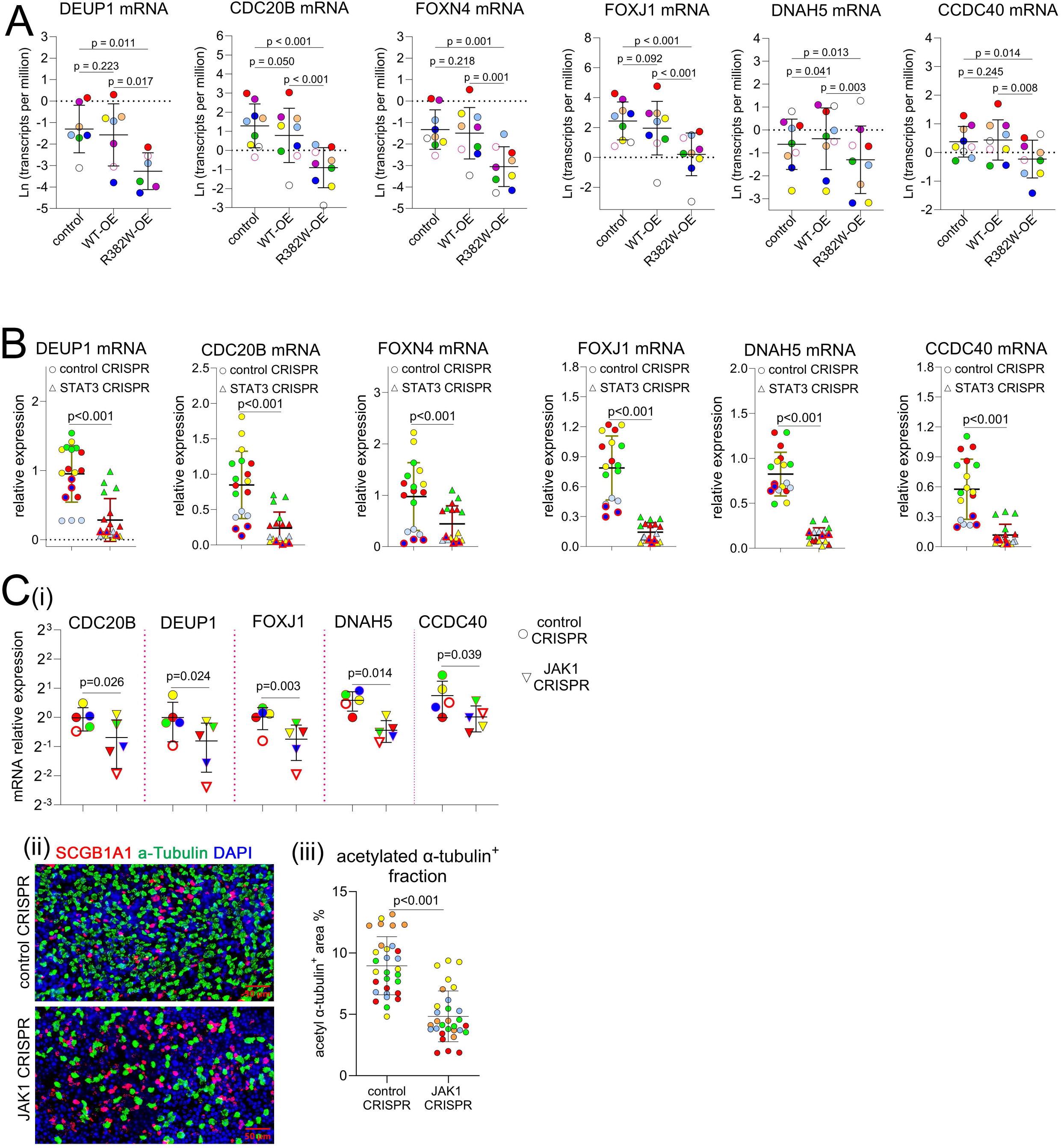
JAK1/STAT3 axis is required for ciliogenesis. **(A)** Expressions of deutrosomal cell markers CDC20B, DEUP1, FOXN4 and multiciliated cell genes FOXJ1, DNAH5, and CCDC40 were measured by bulk RNA-seq in the normal HBE cells (n=9 donors) expressing control, STAT3 WT-OE, and STAT3 R382W-OE lentiviruses and cultured under ALI condition. **(B)** Expressions of **(i)** DEUP1, CDC20B, FOXN4, and **(ii)** FOXJ1, DNAH5, and CCDC40 were measured by TaqMan assays in STAT3-KO (by CRIPSR/Cas9) HBE cells (from 5 donors). **(C)** Expressions of **(i)** CDC20B, DEUP1, FOXJ1, DNAH5, and CCDC40 were measured by TaqMan assays in JAK1-KO (by CRIPSR/Cas9) HBE cells (from 5 donors). Expressions of **(ii)** SCGB1A1 and acetylated α-tubulin were detected by immunofluorescent whole mount staining in the cultures of control and JAK1-KO HBE. **(iii)** The acetylated α-Tubulin signals were quantified by morphometrics. Symbols with the same color in the same panel indicate cultures originated from the same donor. IF: immunofluorescence; HBE: human bronchial epithelial; OE: overexpression; KO: knock-out.

**Supplementary Figure 10:**
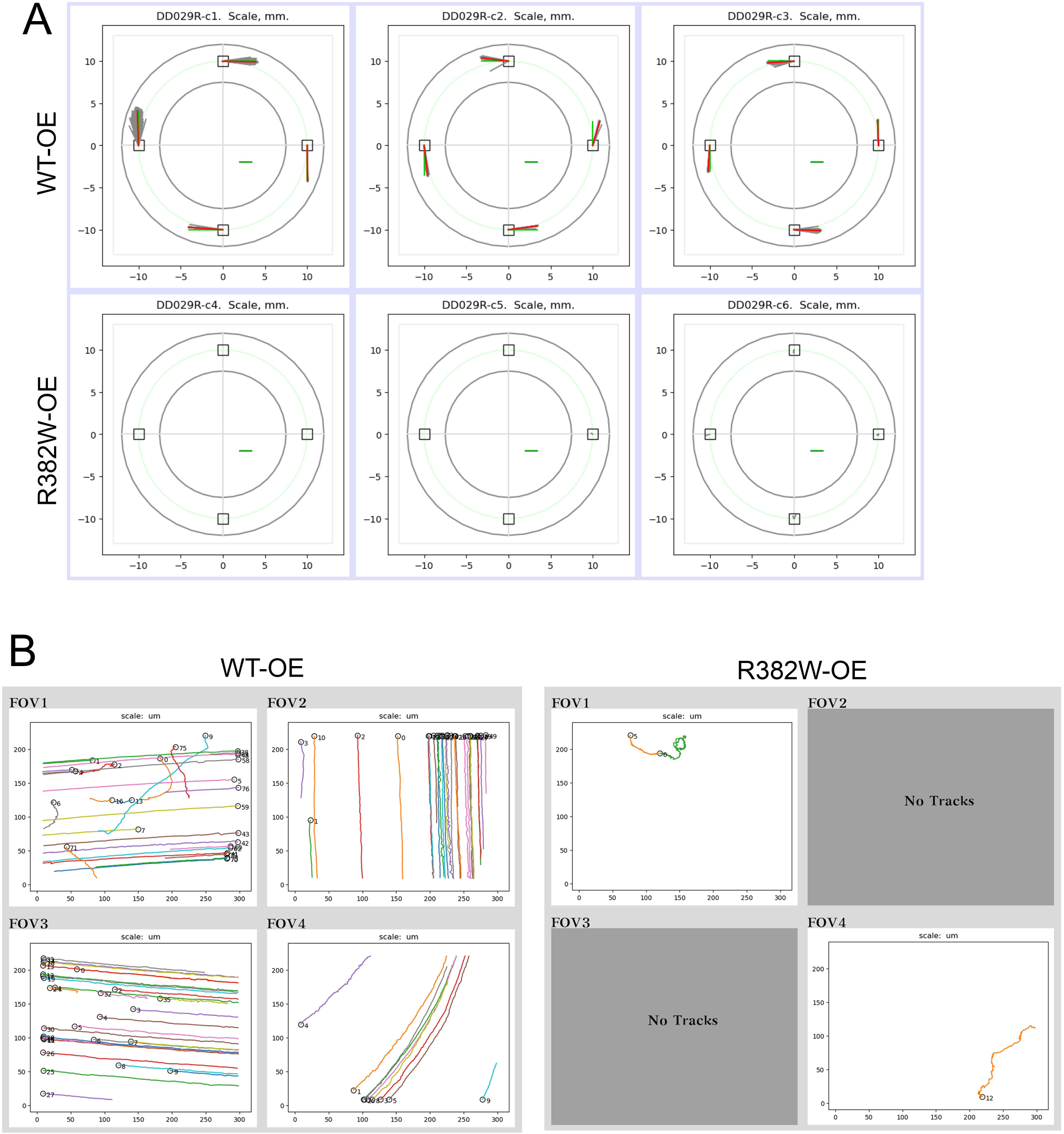
STAT3 R382W mutation leads to abnormal mucociliary transport. **(A)** Representative transport velocity maps of HBE cultures expressing STAT3 WT-OE and STAT3 R382W-OE. The velocities for individual and mean tracks are shown in gray and red, respectively. The velocity for global transport is shown in green. The horizontal green bar in quadrant IV is a scale bar for the vectors, 100 µm/s. **(B)** The track images showed globally coordinated transport in WT-OE in the 4 fields of view (FOV, marked as squares presented in **A**) of each culture, and uncoordinated or undetectable transport in R382W-OE HBE culture. Images are representative of HBE cultures from 8 donors. HBE: human bronchial epithelial; OE: overexpression.

**Supplementary Figure 11:**
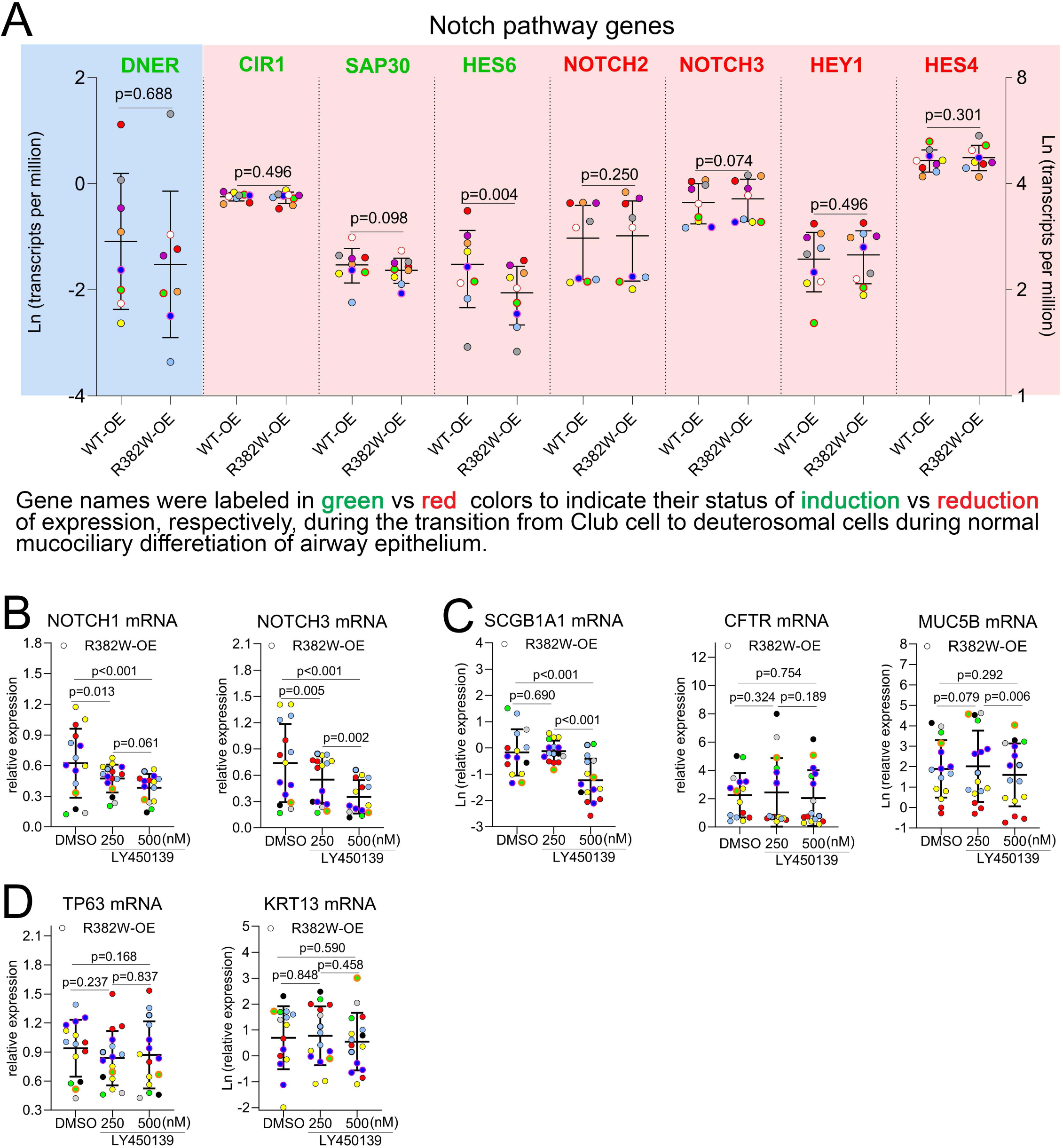
Gamma secretase inhibitor LY450139 (250nM) inhibits NOTCH pathway without changing expression of secretory, basal and suprabasal cell markers in STAT3 R382W-OE HBE cells. **(A)** A group of selective genes involved in the Notch pathway that regulate the transition from Club/secretory cells to deutrosomal/multiciliated cell lineage during normal epithelial cell ciliogenesis (referenced by the published longitudinal airway epithelial cell mucociliary differentiation scRNA-seq data) were compared in STAT3 WT-OE vs STAT3 R382W-OE HBE cells (from 9 donors). Gene names were labeled in green vs red colors to indicate their expression status of induction vs inhibition, respectively, during this transitional period. The data presented in light blue areas were plotted to left y axis and data presented in pink areas were plotted to the right y axis. Expression of **(B)** Notch pathway activators NOTCH1, NOTCH3, **(C)** secretory cell markers SCGB1A1, CFTR, MUC5B, and **(D)** basal cell marker TP63 and suprabasal KRT13 in response to LY450139 treatment (250nM and 500nM) or vehicle (DMSO) was determined by TaqMan assay in STAT3 R382W-OE HBE cells (n=9 donors). Symbols with the same color in the same panel indicate cultures originated from the same donor HBE: human bronchial epithelial; ALI: air-liquid interface; OE: overexpression.

**Supplementary Videos 1-5:**

Normal human bronchial epithelial (HBE) cells expressing STAT3-WT (WT-OE) and STAT3-R382W mutant (R382W-OE) were seeded on the homemade racetrack device and cultured at air-liquid interface conditions for 8 weeks prior to the measurement of surface ciliation (ciliary beating active area), ciliary beat frequency (CBF), and mucociliary transport rate. The bright field video clips demonstrate cilia beating in WT-OE (**Supplementary Video 1**) and R382W-OE (**Supplementary Video 2**) HBE cultures. Fluorescent beads (2µm, carboxylate modified) were added to the cultures, and the movement of these beads, driven by mucociliary transport, was recorded using high-speed video microscopy. The WT-OE HBE cultures exhibited both globally coordinated and globally uncoordinated transport activities, as shown in **Supplementary Videos 3** and **4**, respectively. In contrast, globally coordinated transport was completely absent in R382W-OE HBE cultures. Most of the R382W-OE cultures (6 out of 8 donors) showed no detectable transport, while a few (2 out of 8 donors) displayed uncoordinated transport (**Supplementary Video 5**). These videos represent cultures from n=8 donors.

## Supplementary Tables

**Supplementary Table 1:** Information of people with STAT3-HIES and healthy controls who contributed to this study. STAT3 mutation types and their domains presented in STAT3 protein from people with STAT3-HIES were defined and showed. The age and sex of the donors with disease and healthy controls were also listed in the tables.

| CODE # | Mutation | STAT3 region | Gender | AGE |
| --- | --- | --- | --- | --- |
| STAT3-HIES#1 | 1853 G>A; G618D | SH2 | F | 31 |
| STAT3-HIES#2 | 1027 G>T; V343F | DNA binding | M | 27 |
| STAT3-HIES#3 | 1997T>G; L666R | SH2 | M | 28 |
| STAT3-HIES#4 | 1144C>T; R382W | DNA binding | F | 44 |
| STAT3-HIES#5 | 1144C>T; R382W | DNA binding | M | 32 |
| STAT3-HIES#6 | 1387delGTG; V463del | DNA binding | F | 21 |
| STAT3-HIES#7 | 1144 C>T; R382W | DNA binding | F | 40 |
| STAT3-HIES#8 | 1144C>T; R382W | DNA binding | M | 32 |

**Supplementary Table 1: Information of people with STAT3-HIES and healthy controls who contributed to this study.**
| CODE # | Gender | AGE |
| --- | --- | --- |
| control#1 | F | 31 |
| control#2 | M | 27 |
| control#3 | M | 28 |
| control#4 | F | 44 |
| control#5 | M | 32 |
| control#6 | F | 21 |
| control#7 | F | 40 |
| control#8 | M | 32 |

**Supplementary Table 2:**
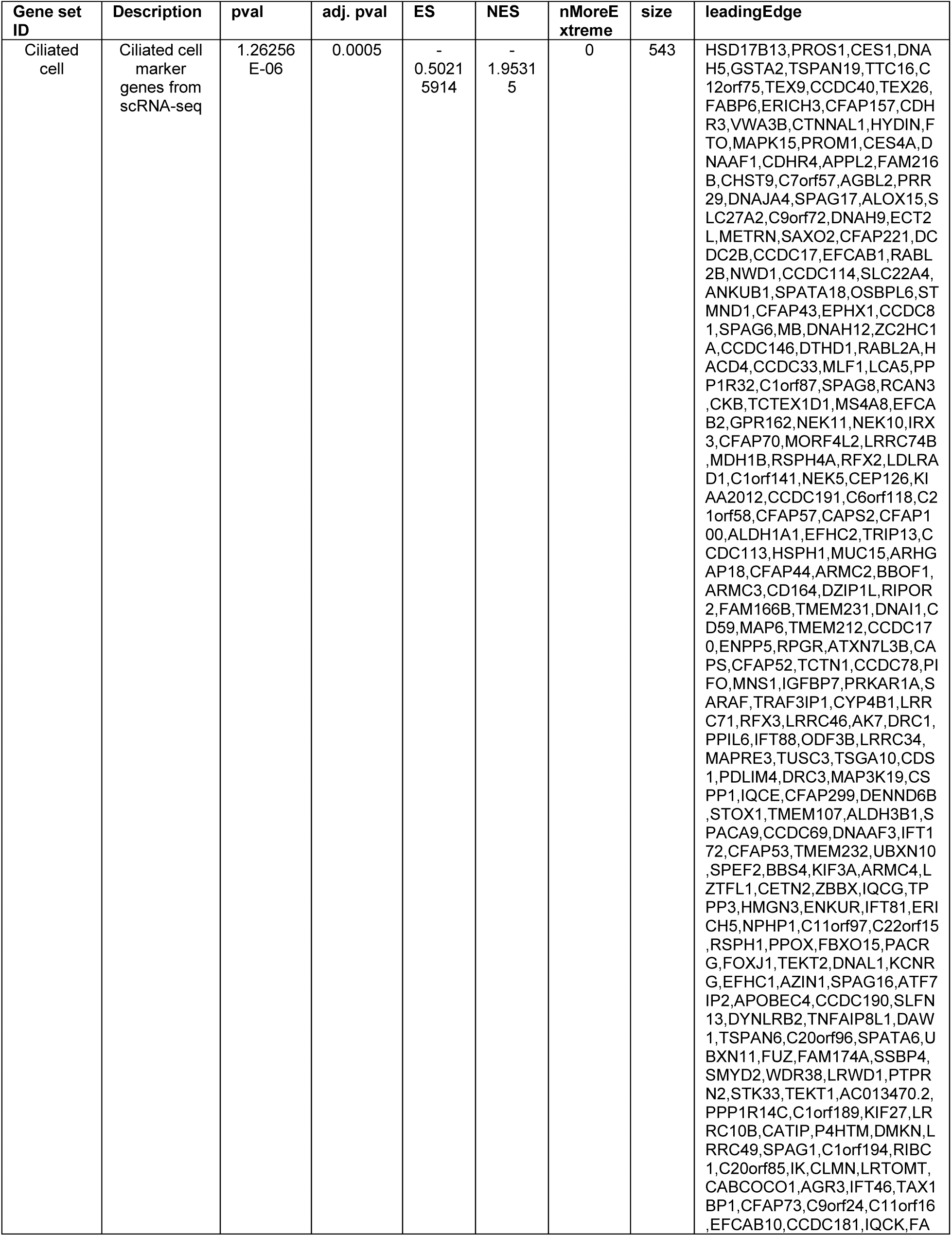
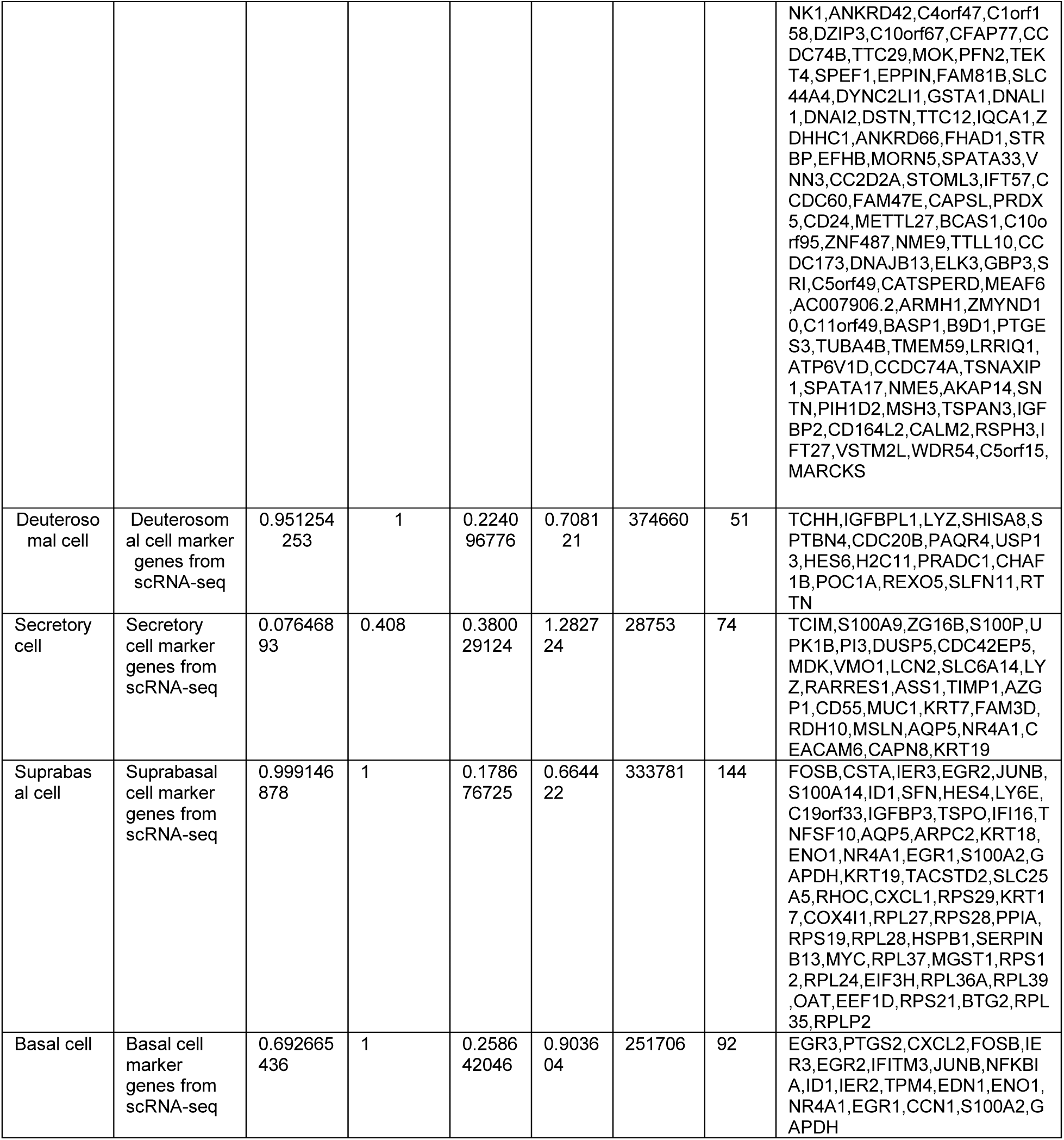
The gene sets selected for generation of ridge plot that predicts cell types in the airway epithelium of STAT3-HIES lung. The relatively enriched gene sets presented individual major airway epithelial cell type were previously identified by scRNA-seq from the biopsy tissues from normal human bronchus (Okuda et al 2021 AJRCCM). The gene sets were then selected to test differential expression in the epithelium of bronchial brushing tissues of STAT3-HIES vs control subjects, and genes that contributed to generate leading edge of the ridge of each cell type were listed, and the number of the genes enriched in the subtype of the epithelium contributed the “size” column. **pval**: nominal p-value of enrichment; **adj. p val**: multiple test corrected p-value; **ES**: enrichment score; **NES**: normalized enrichment score accounting for gene set size; **nMoreExtreme**: Number of times a better ES was achieved throughout number of permutations (typically, 1 million); **leadingEdge**: leading genes that drive the enrichment.

## Materials and Methods

### Human Specimens

Human specimens used in this study were collected under the supervision of Dr. Kenneth Olivier’s team at the National Heart, Lung, and Blood Institute (NHLBI), in accordance with IRB protocol #07-H-0142. Informed consent was obtained from all participants prior to enrollment in the study.

### Sputum Collection

#### Spontaneously Produced Sputum

Participants who were able to spontaneously produce secretions without the influence of nebulizer saline from the sputum induction procedure spit saliva into a cup for discard, then coughed up a sputum specimen into a sterile cup for collection.

#### Induced Sputum

For participants unable to produce a sputum specimen, an induced sputum protocol was applied. The participant used a jet nebulizer device with 4 ml of sterile 7% saline solution for 20 minutes. Saliva was spit into a cup for discard. After spitting, sputum brought up by coughing was collected into a sterile cup. Collected sputum samples were immediately stored at −80°C for later analysis.

### Bronchial Brushing

Bronchial brushing samples were collected by trained pulmonologists using flexible bronchoscopes (Olympus). Prior to the procedure, participants underwent standard preoperative preparations, including fasting and administration of local anesthesia to the oropharynx and larynx. Bronchial brushing was performed by passing a sterile, disposable cytology brush (Olympus Respiratory Endoscopy Cytology Brush) through the working channel of the bronchoscope and gently brushing the surface of the bronchial mucosa with direct visualization. This procedure was repeated 5-10 times and on the contralateral side. Following collection from the participant, one part of the bronchial brushing tissues was immediately preserved in RNAlater for later bulk-RNA-seq, and the other part was transferred to a sterile container with RPMI-1640 supplemented with antibiotics prior to overnight shipping to UNC at Chapel Hill.

### Bronchoalveolar Lavage (BAL)

The bronchoscope was carefully inserted through the mouth, employing appropriate local anesthesia to minimize discomfort. The bronchopulmonary segment for BAL was selected based on predetermined criteria, with emphasis on the middle lobe or lingula. Sterile saline solution was instilled into the target bronchopulmonary segment in aliquots of 50ml through the working channel of the bronchoscope. The instilled saline was then gently aspirated using a suction apparatus, and the lavage fluid was collected into labeled sterile specimen containers. This fluid was stored at −80°C until analyzed.

### STAT3-HIES Autopsy Lung Specimens

Lung sections from STAT3-HIES patients were provided by Dr. Alexandra Freeman of the National Institute of Allergy and Infectious Diseases (NIAID). These excised tissues were processed and embedded in paraffin, then sectioned at a thickness of 5 µm for subsequent analysis.

### Mucus %solids and total, individual mucin measurement

#### Measurement of mucus solids content in sputum samples

The percentage of solid content of mucus, an index of hydration, was calculated by measuring wet/dry weights with a microbalance (UMX2; Mettler Toledo). For measuring the sputum %solids, we followed the protocol previously described (1). The solids concentration in sputum mucus was determined by aliquoting 100µl of sputum onto a pre-weighed piece of foil. The initial mass of the sputum along with the foil was recorded. The sample was then dried in an oven set at 80°C overnight. Following the drying period, the final mass of the dried sample and the foil was recorded. The percentage of solids by weight was then calculated based on these measurements.

#### Measurement of mucus solids content in the apical secretions from human bronchial epithelial cell cultures

To assess the solids content in the apical secretions following cytokine treatment, the filter paper technique previously described (2–4) was employed. STAT3 WT-OE and R382W-OE lentiviruses-infected human bronchial epithelial (HBE) cells were cultured for four weeks under air-liquid interface conditions using UNC-ALI media (5). Cells were then cultured for an additional 5 days prior to treatment with IL1β cytokine at a concentration of 1 ng/ml (added in media) for 3 days. The apical surface remained unwashed during this 8-day period to allow accumulation of secretions. Post-treatment, the apical secretions were collected, and their dry versus wet weights were measured. The percentage (%) of solids by weight was calculated based on these measurements.

#### Total mucin concentration and specific mucin quantification

Total mucin concentrations in the sputum were quantified using size-exclusion chromatography combined with differential refractometry (6). Sputum samples were injected into a CL2B size-exclusion column to isolate mucins. The isolated mucins were then passed through an enhanced optimal system laser photometer (DAWN HELEOS II, Wyatt Technology) coupled with an interferometric refractometer (Optilab T-rEX, Wyatt Technology) for continuous measurement of light scattering and sample concentrations. Data analysis was performed using ASTRA software version 6.1.1.7 (Wyatt Technology), which utilized the ASTRA Pro algorithm for molecular weight distribution calculations.

To specifically measure MUC5AC and MUC5B concentrations, sputum samples were reduced, alkylated, and digested with trypsin (6). Internal standards consisting of six heavy-labeled peptides (three for each mucin) were added to the digested samples at known concentrations. The samples were then fractionated using high-performance liquid chromatography (HPLC) and analyzed by targeted selected-ion monitoring–data-independent acquisition (SIM-DIA) using a Dionex Ultimate 3000 RSLCnano system coupled to a Q-Exactive hybrid quadrupole-orbitrap mass spectrometer with a Nano spray source (Thermo Fisher, Bremen, Germany). Data analysis was performed using Skyline software (MacCross Lab, version 20.1): peak areas for each peptide (endogenous and heavy internal standard peptides) are calculated based on precursor ion (mass spectrometry) and top three highest intensity product ions (tandem mass spectrometry). Ratios from three peptides were averaged and MUC5B and MUC5AC concentrations were calculated.

### Sialic acid and urea concentration measurement

The concentration of sialic acid and urea were measured by mass spectrometry as previously described (7). Briefly, after the addition of isotopically labeled internal standard solution, samples were incubated with 1% formic acid at 80°C for one hour to release sialic acid from mucins, then centrifuged through a 10-kDa size-selection filter (EMD Millipore, Billerica, MA) to remove high molecular weight compounds. 5 µL was analyzed by liquid chromatography-tandem mass spectrometry using a UPLC T3 HSS C18 column with methanol/formic acid gradients for chromatography and a Quantum-ultra triple quadrupole mass spectrometer (ThermoFisher, Waltham, MA) in positive mode. Peak area for each metabolite was determined using automated software (Xcalibur, ThermoFisher), with concentrations assessed using ratios to the relevant internal standard and compared to standard solutions run in parallel.

### Macroscopic rheology

Macroscopic rheology assessments were conducted using a TA Discovery Hybrid Rheometer 3 (TA Instruments, New Castle, Delaware) equipped with a 20mm diameter 10° cone, following a previously described method (8). All experiments were performed at 23°C and included a solvent trap to prevent sample dehydration. The following procedures were applied to each specimen:

#### Stress amplitude sweeps

Stress sweeps were conducted at frequencies of 0.1 and 10 Hz with stress ranging from 0.01 Pa to 1 Pa. This step determined the linear viscoelastic regime (LVR) of the mucus specimens.

#### Frequency sweeps

Frequency sweeps were carried out from 0.01 to 10 Hz at stress levels below the nonlinear threshold to measure macroscopic linear moduli. Rheological data were analyzed using TA Trios software. The complex viscosity (η*) from the amplitude sweeps was directly recorded and processed by the software.

### Sample preparation for Immunohistochemistry, Alcian Blue Periodic Acid Schiff (AB-PAS) staining, and confocal microscopy

Normal lung tissues containing bronchi, bronchioles, and terminal bronchioles were obtained from the Tissue Procurement and Cell Culture Core at UNC Marsico Lung Institute, in accordance with IRB protocol #03-1396. The tissues were fixed in 10% neutral buffered formalin for 48 hours through immersion, followed by paraffin embedding. Sections were then cut to a thickness of 5μm. Concurrently, air-liquid interface (ALI)-cultured human bronchial epithelial (HBE) cells were fixed on the transwell membrane with 10% neutral-buffered formalin for 1 hour at room temperature and washed with PBS prior to embedding and sectioning. Additionally, 100µl of sputum mucus from each donor was applied to slides using a cytospin device and allowed to air dry.

#### Staining procedures

Hematoxylin and eosin (H&E), Alcian Blue Periodic Acid Schiff (AB-PAS) staining, and both immunohistochemical and immunofluorescent staining were performed according to previously described methods (9).

#### Confocal microscopy for air-liquid interface (ALI)-cultured cells

ALI-cultured human bronchial epithelial (HBE) cells prepared for whole mount analysis were mounted on glass slides using a DAPI-containing medium and covered with coverslips. Imaging was conducted using a Leica Stellaris 5 confocal microscope with 63x 1.4 NA oil immersion lenses. Fluorescence excitation was tailored using appropriate laser lines, with detector settings adjusted to optimize signal-to-noise ratios. The pinhole was set to 1 Airy Unit. Z-stacks were captured as necessary using LAS X software, which was also employed to adjust image contrast and brightness. Further analysis was conducted using Qupath software (10).

### U-PLEX Assay for Cytokine and Chemokine Analysis

Cytokine and chemokine concentrations in bronchoalveolar lavage (BAL) and apical lavage or basolateral media from human bronchial epithelial (HBE) cell cultures were quantified using the U-PLEX assay (Meso Scale Diagnostics, Rockville, MD, USA). This assay utilizes an electrochemiluminescent detection method and was performed following the manufacturer’s recommendations.

#### Cell culture and treatment

HBE cells were cultured in Vertex ALI (VALI) media (11) for four weeks. Subsequently, cells were treated with IL1β applied basolaterally at a concentration of 1 ng/ml for 24 hours. Prior to sample collection, 200 µl of PBS was added to the apical side of the cultures and incubated for 15 minutes. Both apical lavages for meso-scale analysis and basolateral media were collected to measure the concentrations of cytokines and chemokines.

#### Assay procedure

On the day of analysis, samples were thawed, and 96-well plates were prepared by coating with linker-coupled capture antibodies for one hour. After aspiration, plates were washed three times with PBS containing 0.05% Tween-20 (PBST). Standards and samples (25 μl) were then added to the appropriate wells and incubated for one hour at room temperature with shaking. After incubation, wells were washed three times with PBST. Detection antibodies were added to each well and incubated for an additional hour at room temperature, followed by another three washes. Each well then received 150 μl of reading buffer. Plates were immediately analyzed using the MESO QUICKPLEX SQ 120 instrument, equipped with DISCOVERY WORKBENCH® data analysis software (Meso Scale Diagnostics). Standard curves were generated by fitting the electrochemiluminescence signals from calibrators to a weighted 4-parameter logistic model.

### Expansion of human bronchial epithelial (HBE) cells from bronchial brushing tissues

#### Collection of bronchial brushing tissues

Bronchial brushing tissues were collected from healthy donors and donors with STAT3-HIES. The brush portion containing the tissues was severed from the cytology brush and placed in RPMI media within 1.5 ml Eppendorf tubes. These samples were stored at 4°C and shipped overnight from the NHLBI in Bethesda, MD, to UNC-Chapel Hill, NC.

#### Tissue processing and cell dissociation

Upon arrival the next morning, tissues were transferred to cell dissociation buffer, consisting of 4.5 ml Accutase and 0.5 ml of freshly prepared 5 mg/ml Pronase (Sigma-Aldrich, cat #SCR005 and #10165921001). The tissues were incubated at room temperature for 1 hour on a 3-D rocker to facilitate dissociation. Dissociation was halted by adding 0.5 ml of fetal bovine serum (FBS). The cell mixture was further dissociated by pipetting up and down 10 times using a 10 ml pipette.

#### Cell harvesting and expansion

After removing the brush tip, the cell suspension was centrifuged at 1000g for 2 minutes to pellet the cells. The supernatant was discarded, and the pellet was resuspended in conditional reprogramming culture (CRC) media (see the Reagents Table-1). Expansion of the isolated bronchial brushing epithelial cells followed the conditional reprogramed cell (CRC) culture protocol previously described (12–14) with minor modifications. The isolated cells were then seeded onto a 10 cm dish containing 3T3-SA feeder cells (ATCC act#CCL-92), which had been treated with mitomycin-C (Sigma-Aldrich, Cat# M4287-5X2MG) to induce growth arrest and prepared the previous night. Once the bronchial epithelial cells reached confluence, they were counted and cryopreserved at −80°C in cell cryopreservation media for future use.

#### Antibiotics and Antifungal reagents

The STAT3-HIES bronchial epithelial cells isolated from brushing tissues were cultured in CRC media supplemented with antibiotics and antifungal agents, following the protocols our group developed for culturing CF cells isolated from chronically infected lungs (15, 16).

### Primary human bronchial epithelial (HBE) cell culture

Normal primary HBE cells (obtained from donors without previously known pulmonary diseases), or control and STAT3-HIES HBE cells expanded via conditional reprogramed cell culture protocol were differentiated following the protocol previously described (5, 17). HBE cells were maintained at an air-liquid interface (ALI) condition. The apical surface was washed with PBS, and UNC-ALI medium (5, 17) was replaced only in the basal compartment two-three times per week, and cells were cultured under ALI conditions for 4 weeks to allow full differentiation. All HBE cells were cultured using UNC-ALI media until the end of the experiments, unless otherwise specified.

### Ussing chamber assays

In vitro short circuit current (Isc) measurements of CFTR Cl^-^ secretion were assessed in the primary control and STAT3-HIES HBE cells, STAT3 CRISPR-targeted and lentiviruses (control, STAT3 WT-OE, R382W-OE) infected HBE cells as previously described (13). Briefly, HBE cells were cultured on snap wells (Corning #3801) under air-liquid interface (ALI) for 4 weeks prior to IL1β exposure (basolaterally, 1ng/ml) using Vertex ALI media (11). The cells were then mounted in Ussing chambers (Physiologic Instruments, San Diego, CA) for electrophysiological measurements. The Ussing chambers were filled with a bilateral solution (5 ml) of Krebs-bicarbonate-Ringer containing: 140mM Na^+^, 120mM Cl^-^, 5.2mM K^+^, 1.2mM Ca^2+^, 1.2mM Mg^2+^, 2.4 mM HPO4^2-^, 0.4mM H_2_PO4^-^, 25mM HCO_3_^-^, and 5mM glucose. The solution was circulated with 95% O_2_, 5% CO_2_ gas, pH 7.4, and maintained at 36°C +/-1°C. To measure cyclic AMP (cAMP) activation of CFTR, forskolin (10μM) was bilaterally applied in the presence of 100μM amiloride (apical) to inhibit ENaC activity. This was followed by inhibition of CFTR-specific current with CFTR inhibitor-172 (10μM, apical). Transepithelial resistance (Ω·cm2) was measured to assess monolayer integrity.

### Western Blot Analysis

Total proteins from HBE cells cultured under ALI conditions and H441 cells lysed in 6M urea with protease and phosphatase inhibitors (ThermoFisher, Cat#78430, 78428) were processed. Cytoplasmic and nuclear proteins were extracted separately using the Nuclear Extract Kit (Active Motif, 40010). Proteins were quantified and equal amounts per sample were separated on NuPAGE 10% Bis-Tris Gels (polyacrylamide) and transferred onto PVDF membranes using the iBlot™ 2 Transfer Stacks kit.

#### Immunoblotting

Membranes were probed using primary antibodies listed in the Reagents Table-2, followed by IRDye 800CW donkey anti-rabbit IgG and IRDye 680LT donkey anti-mouse IgG (LI-COR Biosciences) as secondary antibodies. Protein detection and densitometry were conducted using the Odyssey Infrared Imaging System (LI-COR Biosciences).

#### CFTR protein detection

CFTR western blot followed the protocol previously described (18). Whole-cell lysates from fully differentiated HBE cells (4 weeks of ALI culture) were lysed using NP-40 lysis buffer supplemented with protease inhibitors. The lysates were clarified, and CFTR protein was immunoprecipitated using a rabbit anti-CFTR polyclonal antibody (UNC CFTR-antibody 155) and Protein A/G PLUS-agarose beads (Santa Cruz Biotechnology). The immunoprecipitated complexes were eluted from the beads, separated by SDS-PAGE, and transferred to a membrane for Western blot analysis. CFTR was detected using primary antibodies (CFTR-596 and 217)(18) specific to CFTR or to actin and TBP (TATA box binding protein) as a loading control, followed by appropriate fluorophore-conjugated secondary antibodies. The proteins were visualized with the Sapphire Biomolecular Imager by Azure Biosystems.

#### Mucin detection

For MUC5B and MUC5AC mucin analysis, we followed the protocol previously described (19). Briefly, 100µl PBS was added to the apical surface of HBE cultures to collect secretions, which were transferred to 1.5 ml tubes and adjusted to 250 µl. Mucins were denatured with urea to a final concentration of 6M. Samples (40µl) were run on 1% agarose gels at 80V for 90 minutes and proteins were transferred to nitrocellulose membranes using a vacuum system. MUC5AC (20) and MUC5B (21) proteins were detected using their specific antibodies. Signal detection and densitometry were performed with the Odyssey Infrared Imaging System (LI-COR Biosciences).

### Plasmid and Lentivirus Generation

The method of generation lentiviral-mediated overexpressing vectors and virus production followed the protocol previous described (4).

#### Plasmid construction

To address the need for specific gene expression in primary human bronchial epithelial cells, we generated lentiviruses designed for overexpression studies using the EF1α-promoter driven lentiviral system (22). The SARS-CoV-2-M-2xStrep fragment was excised from the pLVX-EF1alpha-SARS-CoV-2-M-2xStrep-IRES-Puro vector (Plasmid #141386, Addgene) using EcoRI and BamHI enzymes. The sticky ends of the vector backbone were then converted to blunt ends using DNA Polymerase I, large (Klenow) fragment (NEB, cat# M0210S), and subsequently ligated using T4 DNA ligase (NEB, cat# M0202S). Human STAT3 WT (NM_001369512.1) and STAT3 R382W (C1144T) fragments, synthesized into pcDNA3.1+ plasmids by Genescript Inc., were amplified by PCR and sub-cloned into the pLVX-EF1alpha-IRES-Puro vector through blunt-end ligation.

#### Viral production and titration

To produce high-titer lentiviral vectors necessary for efficient transduction, Lenti-X™ 293T cells (Takara Bio, cat# 632180) were cultured to approximately 80% confluence in 150×25 mm dishes (Sigma-Aldrich, cat# CLS430599-60EA). The cells were transfected with 10μg of transfer plasmid, 5μg of pCMV-VSV-G, 8μg of psPAX2, and 35μl of transfection reagent (Sigma-Aldrich, cat# 6366546001) in Opti-MEM (Thermo Fisher, cat# 51985091. After 6 hours, the media was replaced with D10 media (Thermofisher DMEM, 10% fetal bovine serum, 1% bovine serum albumin; cat# 11965092, #26140079; Sigma-Aldrich, cat# A9418-50G). Viral supernatants were collected 60 hours post-transfection, centrifuged at 4,000 rpm for 10 minutes at 4°C, and filtered through a 0.45μm low protein binding membrane (Millipore, cat# SLHP033RS). Viral titers were determined using a Lenti-X™ GoStix™ Plus kit (cat# 631281, Takara Bio) according to the manufacturer’s instructions.

### Lentivirus Infection

#### Primary human bronchial epithelial (HBE) cells

Primary HBE cells were cultured on 10 cm dishes coated with PureCol (Advanced BioMatrix, 5005-B) in modified CRC media. We seeded approximately one million passage 1 (P1) cells per dish, allowing them to achieve 30-50% confluency. Lentiviruses were administered at a multiplicity of infection (MOI) of 1 together with 1µg/ml of polybrene (Sigma-Aldrich, Cat#TR-1003-G) in CRC media, and the cells were incubated for 4 hours before the viral media was removed. Following infection, the cells were maintained in CRC media for 48 hours. Subsequently, they were passaged onto 15 cm dishes, also coated with PureCol, and subjected to puromycin (ThermoFisher, Cat#A1113802) selection (1μg/ml in CRC media). After reaching confluence, the cells were either cryopreserved in liquid nitrogen or transferred to Corning Transwells containing CRC media supplemented with 1μg/ml puromycin. Once confluent, cells were cultured under air-liquid interface (ALI) conditions using UNC-ALI media with 0.5 μg/ml puromycin until the termination of the culture.

#### H441 cell line

The infection protocol for the H441 cell line followed the same procedures, except that the culture medium used was RPMI 1640 supplemented with 10% fetal bovine serum (FBS) and penicillin-streptomycin instead of CRC media.

### Electroporation of CRISPR/Cas9

The control, STAT3, JAK1 and IL1R1(23) sgRNAs, and Cas9 protein were ordered from Synthego. Ribonucleoprotein (RNP) complexes were prepared by combining 360 pmol of single-guide RNA (sgRNA) with 40 pmol of Cas9 protein in 50 µL of Buffer R. The mixture was incubated at room temperature for 10 minutes and subsequently chilled on ice. Human bronchial epithelial (HBE) cells, numbering 0.3 million, were suspended in 200 µL of Buffer R and mixed with the RNP complex. Electroporation was conducted immediately using a Neon Transfection System (Thermo Fisher Scientific) with the settings adjusted to 1600V, 20 msec, and a single pulse. Following electroporation, cells were cultured for expansion and ALI cultures.

*PCR amplification and sequencing analysis:* DNA was extracted from well-differentiated cells post-expansion. PCR amplification of the target region was performed using specifically designed primers. The resulting PCR products were sent for Sanger sequencing. Analysis of the knockout efficiency of the target gene was performed using the ICE (Inference of CRISPR Edits) tool available on the Synthego website.

### IL6 concentration measurement from IL1β treated HBE cells

Well-differentiated human bronchial epithelial (HBE) cells were treated with IL1β at a concentration of 1 ng/ml at four time points: 0 hours, 8 hours, 24 hours, and 72 hours. Following each treatment interval, the apical surface of each culture was lavaged with 200 µl of cell culture grade PBS (Sigma-Aldrich). Both the apical lavages and the basal media were collected. The collected samples were diluted in a 1:9 ratio with H_2_O. The IL6 concentration in the diluted samples was determined using an IL6 Human ELISA kit (R&D Systems, HS600C) according to the manufacturer’s instructions.

### Administration of cytokines, pharmacological inhibitors, monoclonal antibody

HBE cells were cultured under air-liquid interface (ALI) conditions for four weeks to promote full differentiation. Throughout this period, the apical surface of the cultures was washed with PBS, and the UNC-ALI media on the basolateral side was refreshed every three days. After four weeks of ALI culture, recombinant human cytokines were administered. Inhibitor treatments were applied 12 hours before cytokine administration, details of which are provided in the Reagents Table-3. Specifically, cytokines IL1β and IL6 were added to the basolateral side of the ALI cultures at a final concentration of 1 ng/ml. Inhibitors were applied to both the apical and basolateral sides, except where noted otherwise. SC144 and Tofacitinib were initially dissolved in DMSO and then diluted with PBS to a volume of 100µl at the required concentration for apical application. Tocilizumab (24) was dissolved directly in PBS and applied to the apical surface, where it remained for one hour before aspiration.

### Whole-mount immunofluorescence staining and quantification

For histological analysis, HBE cells were cultured under ALI conditions for 4 weeks to allow fully differentiation. HBE cultures were fixed with 10% neutral buffered formalin together with the transwell membranes for one hour at room temperature, followed by 3x PBS (10min/time) wash. The membrane was then excised from the transwell apparatus and washed with PBST (0.3% Triton X-100 in PBS) 1 time for 10 minutes to permeabilize and prepare the tissue for antibody binding. The cultures were blocked using Histo-One blocking buffer (Nacalai USA, Cat#06349-64) for 30 minutes to prevent non-specific antibody binding. Overnight incubation with primary antibodies was performed at 4°C, followed by three 5-minute washes with PBST. Secondary antibodies were then applied at room temperature for one hour. Following another series of three 5-minute PBST washes, the membranes were mounted on glass slides using a mounting medium containing DAPI to stain the nuclei.

#### Imaging and analysis setup

Whole-mount slides stained with Monoclonal Anti-Acetylated Tubulin (Sigma Aldrich, T7451) and CCSP (Abcam, ab51673) were scanned using the z-stack function on an Olympus VS200 slide scanner. The acquired z-stack planes were condensed into a single layer using maximum intensity projection. Each membrane image was divided into six equal sections using the crop function in OlyVIA software. This segmentation aimed to ensure optimal focus level and signal-to-noise ratio presentation of each channel by adjusting the z-axis, as threshold values for highlighting positive areas varied throughout the membrane.

#### Quantification of cilia coverage

Percentage of area covered with ciliated cells was determined from membrane culture scans labeled for acetylated tubulin. An ROI was selected which excluded edges of the culture membrane. Tubulin label regions were thresholded and expressed as a percentage of the total ROI area for six cultures.

#### Secretory cell quantification

CCSP Ab staining was used to quantify the number of secretory cells in the same samples as above. CCSP positive cells were thresholded. The total number of positive cells was calculated by establishing a size (in µm²) representative of one cell, which was used to allow for the division of clusters in order estimate the number of individual cells.

### SEM (Scanning Electron Microscopy)

Primary control and STAT3-HIES HBE cells were cultured under air-liquid interface conditions for 6 weeks using PneumaCult ALI media (Stem Cell, Cat#05001) to promote mucociliary differentiation. Cells were then fixed with 10% NBF (neutral buffer formalin) for paraffin blocks followed by histological assessment by H&E staining. Parallelly, HBE cells were processed to SEM specimens. SEM samples were fixed in 2.5% glutaraldehyde in 0.1M sodium cacodylate buffer. After fixation, samples underwent osmication and dehydration processes. Samples were dried using a Samdri-795 liquid CO2 critical point drier (Tousimis Research Corp, Rockville, MD). Once dried, they were mounted on aluminum planchets using carbon adhesive tabs and coated with a 5 nm thick gold-palladium alloy (60Au:40Pd) utilizing a Ted Pella 208HR sputter coater (Ted Pella Inc, Redding, CA). Imaging was performed on a Supra 25 field emission scanning electron microscope (Carl Zeiss Microscopy, Peabody, MA) at a 5 kV acceleration voltage, with a 5 mm working distance and a 30 µm aperture (25, 26).

### Measurement of Airway Surface Liquid (ASL) pH

The pH measurement of the ASL was conducted based on a modified version of a previously described protocol (27). Human bronchial epithelial (HBE) cells were transduced with lentiviruses expressing either wild type STAT3 (WT-OE) or a STAT3 mutant (R382W-OE) and cultured at an air-liquid interface (ALI) for four weeks to promote differentiation. Post-differentiation, the cells were treated from the basolateral side with either vehicle (PBS) or interleukin-1β (IL-1β, 1 ng/ml) for 24 hours, without washing, to maintain the integrity of the mucus layer.

For ASL pH measurement, 120 µL of Krebs-Ringer bicarbonate buffer (Sigma-Aldrich, Cat# K4002-10x1L) was applied to the apical surface and allowed to equilibrate for four hours at 37°C in an environment of 95% humidity and 5% CO₂. The Orion™ 9810BN Micro pH Electrode was calibrated at room temperature using standard pH buffers (pH 4.01, 6.86, and 9.18) prior to measurements.

The transwell plates with HBE cells were placed on a lab jack inside of the cell culture incubator, allowing manual height adjustment to position the tip of the micro-pH probe approximately 1mm submerging into the apical buffer without contacting the cell layer. A temperature probe continuously monitored the incubator’s internal conditions. Both the micro-pH probe and a temperature probe were connected to a pH meter positioned outside the incubator, with thin cables passing through the rubber seal of the incubator’s glass door to maintain environmental integrity. pH readings were taken 5-6 minutes after submerging the probe into the ASL to allow stabilization of temperature and CO₂ levels at 37°C and 5%, respectively. Following each measurement, the plate was lowered, repositioned to the next transwell, and carefully adjusted to the same height to maintain consistent measurement conditions across all samples.

### Total RNA extraction, reverse transcription, Taqman assays and quantitative RT-PCR

Bronchial brushing tissues from healthy control and STAT3-HIES donors were collected after brushing and immediately preserved in RNAlater (ThermoFisher, Cat#AM7020) until lysis with Trizol followed by RNA purification using Direct-zol RNA Miniprep purification kit (Zymo Research, R2051). HBE cells grown on transwell membranes were also processed similarly. The membrane with cells was excised from the transwell apparatus and lysed in Trizol at 37°C on a shaker at 220 rpm for 30 minutes. Total RNA was purified from these lysates using the Direct-Zol RNA Kit. RNA quantity and quality were assessed using a NanoDrop One Spectrophotometer (ThermoFisher). Subsequently, 200 ng of total RNA was reverse transcribed into cDNA using a Verso cDNA Kit (ThermoFisher, cat# AB-1453/B) at 46°C for 1 hour.

#### Quantitative RT-PCR

The resulting cDNA was diluted 1:20 by molecular grade water for RT-PCR analysis. Quantitative RT-PCR was performed using TaqMan probes (Applied BioSystems) and SsoAdvanced Universal Probes Supermix or ssoadvanced universal SYBR Green supermix (Bio-Rad cat# 1725285, 1725275) on a QuantStudio 6 Real-Time PCR machine (Applied Biosystems). Details of the probes and primers used are listed in the Reagents Table-4.

### Bulk RNA-seq and gene expression analysis

Library Preparation and Sequencing: RNA sequencing libraries were prepared from 1 µg of total RNA using the TruSeq Stranded mRNA Sample Prep Kit (Illumina). mRNA was purified from total RNA using poly-T oligo-attached magnetic beads followed by fragmentation. First-strand cDNA was synthesized using random primers, followed by second-strand cDNA synthesis. The cDNA fragments were then end-repaired, A-tailed, and ligated with indexed adapters. Libraries were enriched with PCR to create the final cDNA library. The quality and quantity of the libraries were validated using the Qubit 2.0 Fluorometer (Life Technologies) and the Agilent 2100 Bioanalyzer. The libraries were sequenced on an Illumina NovaSeq 6000 platform.

#### Data processing and analysis

Raw reads in FASTQ format were aligned to the human reference genome (GRCh38) annotated with current Gencode gene and transcript annotation using STAR (v2.7.3a) (28). Gene expression levels were estimated using StringTie (29) as counts and normalized to Transcripts Per Million mapped reads (TPM).

#### Differential expression analysis

Expression counts were normalized using the ‘voom’ function of the Bioconductor ‘*limma*’ package, which applies mean-variance modeling. Differential expression analysis was conducted with linear mixed-effect models using the ‘dream’ function from the Bioconductor ‘*variancePartition*’ package, treating donor code as a random effect factor. Genes with an adjusted p-value less than 0.05 and an absolute log2 fold change greater than 1 were considered significantly differentially expressed. Principal component analysis (PCA) and heatmap visualization genes from TPM were performed to assess sample clustering and expression patterns using the R packages, *ade4*, *ggplot2* and *ComplexHeatmap*, respectively.

#### Pathway and Enrichment Analysis

Gene set enrichment analysis (GSEA) was performed using the *fgsea* package in R. Pathway gene sets were downloaded from the Gene Ontology (https://geneontology.org/docs/go-citation-policy/), and Reactome (30) databases, while cell type specific markers were derived from human airway epithelial single-cell RNA-seq data set (31). Pathways with an adjusted p-value less than 0.05 were considered significantly enriched.

### Measurements of ciliary beat frequency (CBF), ciliary beating active area, and mucociliary transport (MCT)

To measure ciliated cell activities, STAT3 WT-OE and R382W-OE lentivirus infected HBE cells were seeded on the homemade racetrack device and cultured under ALI conditions for 6 weeks using UNC-ALI: PneumaCult ALI (Stem Cell, Cat#05001) (1:1) mixed media. CBF, ciliary active area (CAA), and MCT measurements were measured following a previously described protocol (32, 33). Following removal of debris with PBS, differentiated HBE cultures were placed within an OkoLab microscope incubator set at 37°C, with humidified air and 5% CO2 flowing at 400 ml/min. Ciliary activity was recorded across 10-12 equally spaced fields on each culture surface using a Nikon Eclipse TE2000 microscope (Nikon Instruments Inc., Melville, NY, USA) equipped with a Basler acA1300-200 um camera. The camera and image capture were controlled using SAVA software (version 2.0.8W, Ammons Engineering, Clio, MI, USA). Videos captured were analyzed with SAVA whole field analysis software to quantify CBF and the extent of CAA.

#### Mucociliary transport measurement

For MCT analysis, fluorescent beads (2µm) were incorporated into the mucus solutions to enhance visualization. The movement of these beads was captured in video format and subsequently analyzed using TrackPy to calculate the average transport speed of mucociliary activity.

### Administration of LY450139 compound

Normal HBE cells infected with STAT3 R382W-OE lentivirus were cultured under ALI conditions using UNC-ALI media for three weeks. After this initial period, these cells were treated with two concentrations of LY450139 (250 nm and 500 nm) or a vehicle control (DMSO), which was added to the basolateral media. The media containing LY450139 was refreshed three times per week. Before harvesting the cells for RNA extraction, ciliary beat frequency (CBF) and ciliary beating active area were assessed. Cells designated for histological analysis were fixed in 10% neutral buffered formalin and prepared for wholemount staining.

### Statistics

#### non-repeated data analysis

Analysis of non-repeated measurement data was performed using Prism 9 software (GraphPad, La Jolla, CA). For comparisons between two groups, Student’s t-test (2-tailed, paired, or unpaired) was used. For comparisons among three or more groups, one-way ANOVA (either matched or not matched) followed by multiple comparison tests (correct for multiple comparisons by controlling the False Discovery Rate) were employed. P values of <0.05 were considered statistically significant. Data are presented as mean ± SD in scatter plots unless otherwise indicated in the figure legends. Non-repeated bulk RNA-seq data were transformed to Ln (natural logarithms) values prior to one-Way ANOVA analysis.

#### Repeated data analysis

To analyze data from repeated measurements of multiple samples from individual sources, a linear mixed-effects model with random intercepts for individual sources and fixed effects for relevant variables was employed. These models were estimated using the PROC MIXED procedure in SAS version 9.4 (SAS Institute Inc., Cary, NC, USA). Wald-tests were utilized to assess the significance of the fixed effects of interest, and the denominator degrees of freedom for these tests were estimated using the Kenward-Roger approximation to accommodate small sample sizes.

#### Data processing prior to linear mixed-effect analysis

Prior to conducting parametric statistical tests, preliminary data analysis included assessments of normality and variance homogeneity. Data exhibiting significant skewness or non-constant variance were considered for transformation to meet the assumptions required by these tests. Logarithmic (log_e_, or Ln) transformation was commonly applied where appropriate to stabilize variance and normalize the distribution, thereby ensuring the robustness and validity of the statistical inferences. The decision to use raw or transformed data was based on these initial assessments, providing a methodological basis for enhancing the analytical accuracy of our statistical approach.

#### Linear regression analysis

The mRNA expression data utilized in this study were expressed as transcripts per million and derived from previously published bulk RNA sequencing datasets. These datasets encompass normal human bronchial epithelial cells cultured in vitro under air-liquid interface conditions until fully differentiated. The references for these datasets are included in previous publications from our group (4, 34, 35). This analysis was carried out using GraphPad Prism software. The linear model was fitted to the data to evaluate the predictive value of the independent variable(s) on the dependent variable, e.g. dynein genes expression on STAT3 mRNA. Key assumptions of the regression, such as linearity, homoscedasticity, independence of residuals, and normality were assessed via the software’s diagnostic tools. The significance of the regression coefficients was evaluated at an alpha level of 0.05. The strength and appropriateness of the model were quantified using the coefficient of determination, r, which reflects the proportion of variance in the dependent variable explained by the independent variables.

### Reagents Table-1 (CRC media)

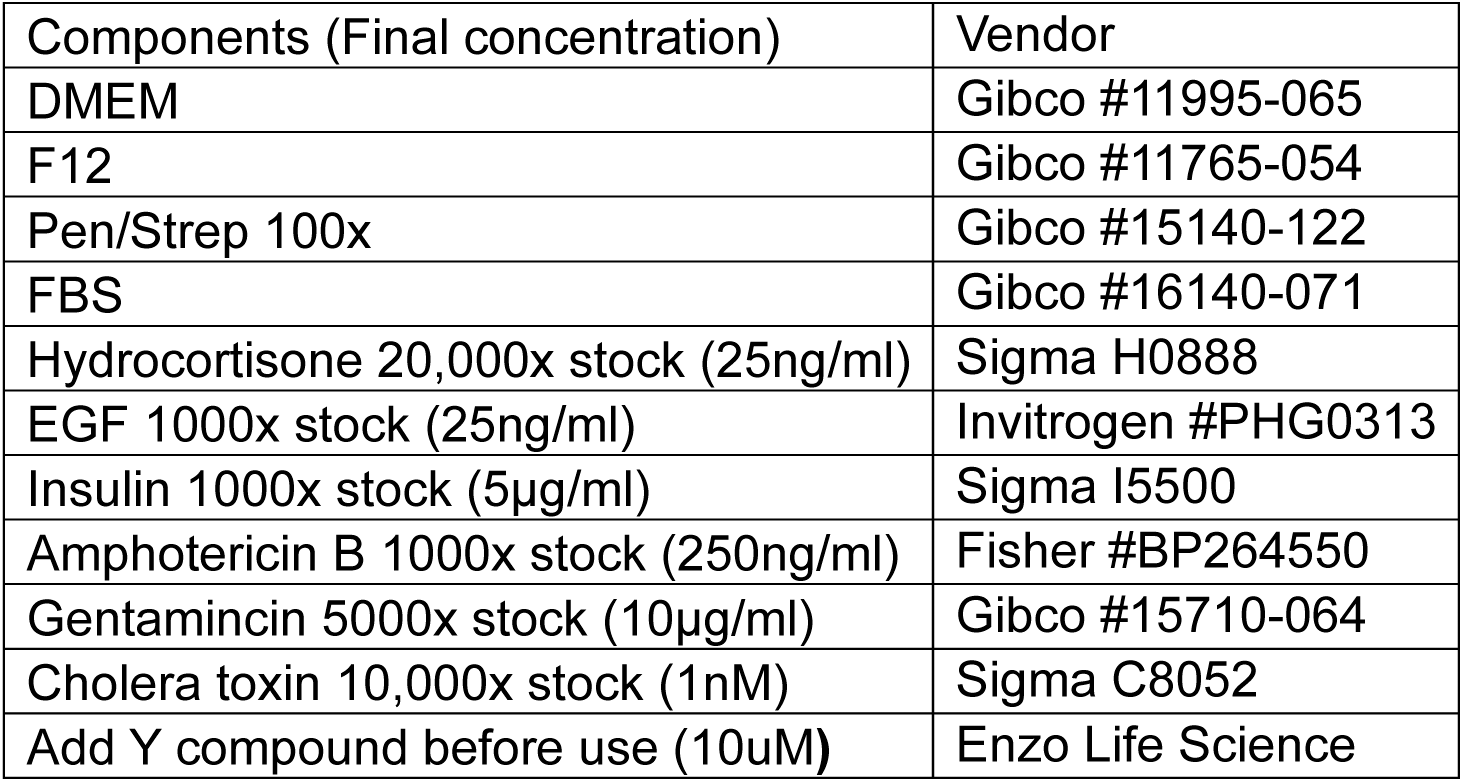

### Reagents Table-2 (Antibodies)

**Reagents Table-2 (Antibodies):**
| Name | Vendor | Cat# |
| --- | --- | --- |
| MUC5AC | Abcam | ab3649 |
| MUC5B | Santa Cruz Biotechnology | sc-20119 |
| IL1B | R&D Systems | MAB201 |
| STAT3 | Cell Signaling Technology | 9139S |
| pSTAT3 | Cell Signaling Technology | 9145S |
| CFTR | UNC CFTR Functional Analysis Core | 596 and 217 |
| TBP | Cell Signaling Technology | 44059S |
| JAK1 | Cell Signaling Technology | 50996S |
| Acetylated α-Tubulin | Sigma-Aldrich | T7451 |
| SCGB1A1 | Sigma-Aldrich | 07-623 |
| β-Actin | Cell Signaling Technology | 4970 |
| GAPDH | Santa Cruz Biotechnology | sc-47724 |

### Reagents Table-3 (recombinant human cytokines and pharmacological inhibitors)

**Reagents Table-3 (recombinant human cytokines and pharmacological** **inhibitors):**
| Name | Vendor | Cat# | final concentration in media |
| --- | --- | --- | --- |
| SC144 | Selleckchem | S7124 | 1 or 10µM |
| Tocilizumab | UNC pharmacy |  | 100µg/ml |
| Tofacitinib | Selleckchem | S5001 | 0.1-10µM |
| IL1b | R&D | 201-LB | 1 ng/ml |
| IL6 | R&D | 206-IL | 1 ng/ml |
| LY450139 | Selleckchem | S1594 | 250nM or 500nM |

### Reagents Table-4 (Taqman and SYBR-Green probes)

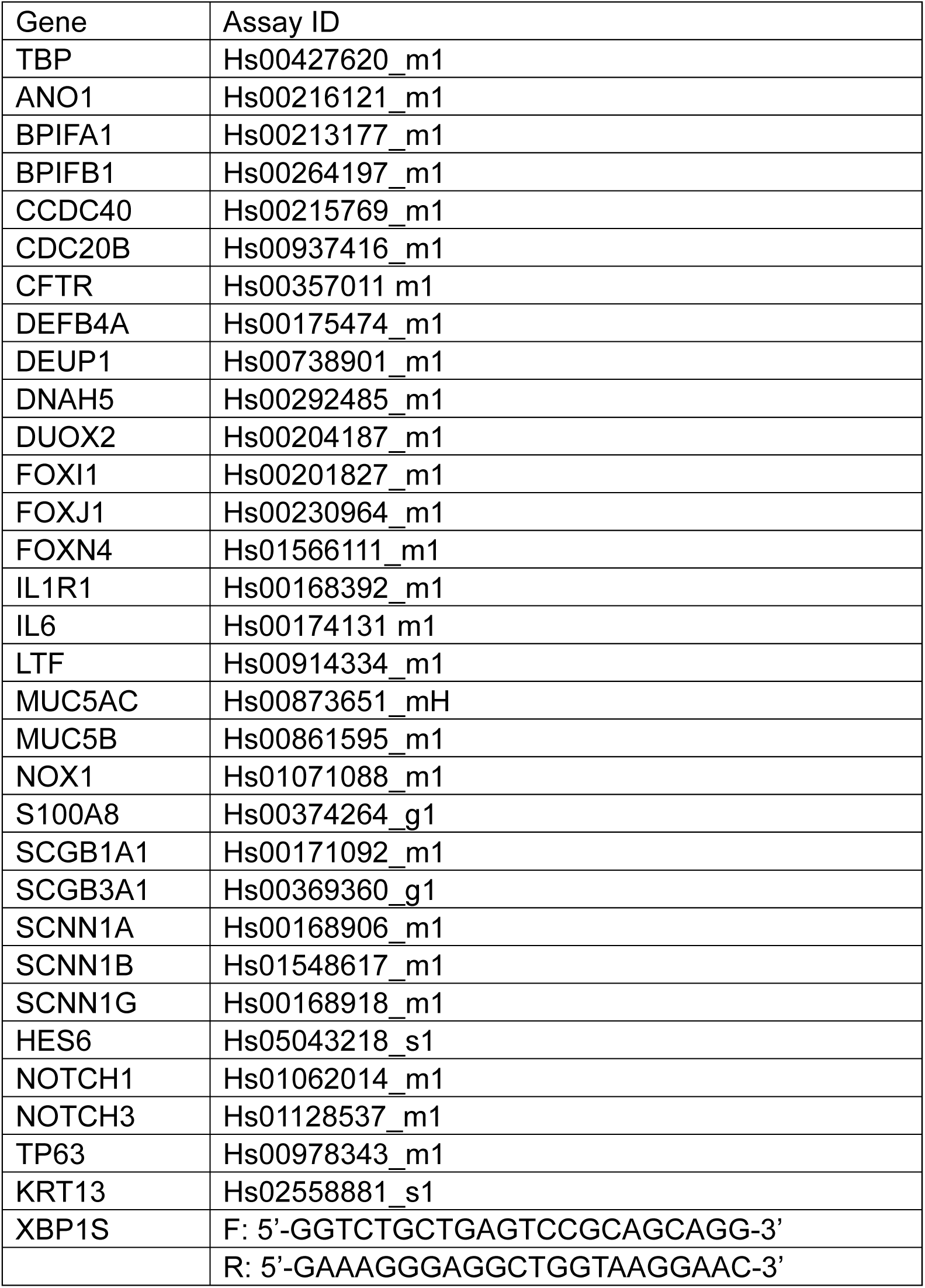

